# Targeting the intestinal circadian clock by meal timing ameliorates gastrointestinal inflammation

**DOI:** 10.1101/2023.01.24.525433

**Authors:** Yunhui Niu, Marjolein Heddes, Baraa Altaha, Michael Birkner, Karin Kleigrewe, Chen Meng, Dirk Haller, Silke Kiessling

**Affiliations:** ZIEL - Institute for Food & Health, Technical University of Munich, 85354 Freising, Germany; Chair of Nutrition and Immunology, School of Life Sciences, Technical University of Munich, Gregor-Mendel-Str. 2, 85354 Freising, Germany; Bavarian Center for Biomolecular Mass Spectrometry, Technical University of Munich, Gregor-Mendel-Str. 4, 85354 Freising, Germany; Faculty of Health and Biomedical Science, University of Surrey, Stagg Hill Campus, GU27XH, Guildford, UK

**Keywords:** circadian clock, colon, gastrointestinal inflammation, IBD, T cells, restricted feeding, microbiota

## Abstract

**Objective:** Impaired clock genes expression has been observed in biopsy samples from patients with inflammatory bowel disease (IBD). Disruption of circadian rhythms, which occurs in shift workers, has been linked to an increased risk of gastrointestinal diseases, including IBD. The intestinal clock balances gastrointestinal homeostasis by regulating the microbiome. Here we characterize intestinal immune functions in mice lacking the intestinal clock and IBD-relevant mouse model under different feeding conditions to describe the functional impact of the intestinal clock in the development of gastrointestinal inflammation.

**Design:** Tissues and fecal samples from intestinal clock-deficient mice (*Bmal1*^IEC-/-^) and mouse models for colitis (*IL-10*^-/-^, *Bmal1*^IEC-/-^x*IL-10*^-/-^, dextran sulfate sodium (DSS) administration) under ad libitum and restricted feeding (RF) conditions were used to determine the causal role of the intestinal clock for colitis.

**Results:** In *IL-10*^-/-^ mice, inflammation correlated with disrupted colon clock genes expression. Genetic loss of intestinal clock functions promoted DSS and IBD inflammatory phenotypes and dramatically reduces survival, and colonization with disease-associated microbiota in germ-free *Bmal1*^IEC-/-^ hosts increased their inflammatory responses, demonstrating the causal role of colonic clock disruption and the severity of IBD. RF in *IL-10*^-/-^ mice restored the colon clock and related immune functions, improved the inflammatory responses and rescued the histopathological phenotype. In contrast, RF failed to improve IBD symptoms in *Bmal1*^IEC-/-^ x*IL-10*^-/-^ demonstrating the significance of the colonic clock to gate the effect of RF.

**Conclusion:** We provide evidence that inflammation-associated intestinal clock dysfunction triggers host immune imbalance and promotes the development and progression of IBD-like colitis. Enhancing intestinal clock function by RF modulates the pathogenesis of IBD and thus could become a novel strategy to ameliorate the symptoms in IBD patients.

## Introduction

The circadian (lat. circa = approximately, dies = day) system consists of the central pacemaker in the suprachiasmatic nucleus (SCN), which drives behavioral rhythms^1^, and orchestrates peripheral clocks that rhythmically regulate tissue-specific functions^2^. Disruption of the circadian system, for example as experienced during jet lag and shift work^3^, relates to immune deficits and pathologies such as cancer, infection^4, 5^, metabolic alterations^6, 7^, and has been associated to a wide range of diseases including inflammatory bowel disease (IBD)^8^.

As previously shown for organs such as the liver, pancreas, muscle and the adrenal, tissue-specific functions are controlled by local circadian clocks^3, 9–11^. Importantly, circadian rhythms have already been identified along the intestine including regeneration of intestinal stem cells, gut motility, nutrient absorption, mucosal immunity and the microbiome^12–17^. Disturbance of GI-physiology results in changes of microbiota composition^18^ and altered microbiota composition plays a role in the development of GI diseases^19^ including IBD^20^. Our recent work on intestine-specific clock deficient mice highlights the importance of functional intestinal clocks for the host-microbe crosstalk in order to balance the host’s GI (GI) homeostasis^21^. Hence, a local oscillator in the intestine may have a strong impact on the development of GI disorders. Indeed, epidemiological studies report suppressed clock genes expression in biopsies from IBD patients^22–24^, indicating a link between the intestinal clock and disease occurrence. However, the importance of the intestinal clock in GI diseases remains to be addressed. Here, we hypothesise that the intestinal clock may constitute a mechanism by which mucosal immune functions and the microbiome promotes GI inflammation and may therefore represent a target for future therapies.

In this study, we provide clear evidence that *IL-10*^-/-^ mice harbours a disrupted colon clock and rhythmicity of various intestinal immune functions and the microbiome is abolished. Importantly, night-time restricted feeding (RF) in *IL-10*^-/-^ mice was identified as a treatment to restore colonic clock functions, induce microbial rhythms, ameliorate the colitis phenotype and enhance their overall survival. Microbiota transfer experiments into germ-free (GF) mice lacking the core clock gene *Bmal1* in the intestine (*Bmal1*^IEC-/-^) directly indicate the causal effect of intestinal clock dysfunction on GI immune responses. RNAseq analysis on colon of *Bmal1*^IEC-/-^ mice validated that key genes relevant for inflammatory and bacterial responses are intestinal-clock controlled. The cause-effect relationship between intestinal clock dysfunction and the severity of GI-inflammation was demonstrated by chemically- and genetically-induced colitis. Remarkably, RF failed to ameliorate the colitis symptoms and survival in *Bmal1*^IEC-/-^ x*IL-10*^-/-^ mice, demonstrating the relevance of a functional intestinal clock for the beneficial effect of RF for IBD.

## Results

### Circadian disruption in an IBD-relevant mouse model

The development of various GI diseases, including IBD^8^ has been linked to circadian disruption, such as occurs during shift work and frequent travel across time zones^3^. In order to characterize whether *IL-10-*deficient mice on Sv129 background (*IL-10^-/-^* ^Sv129^), a well-described mouse model for colitis^25^, exhibit a circadian phenotype, 20 weeks old male mice were kept in different light and feeding conditions (**Fig. 1A, Suppl. Fig. 1A**). Central circadian clock functions, such as rhythmicity of wheel running activity and night time activity during a rhythmic 12-hour light/12-hour dark (LD) cycle and for at least 14 days in constant darkness (DD) was undistinguishable from controls, although total activity was slightly reduced in *IL-10^-/- Sv129^* mice (**Fig. 1A, Suppl. Fig. 1B, C**). No difference in food intake was identified (**Suppl. Fig. 1D**).

**Figure 1.**
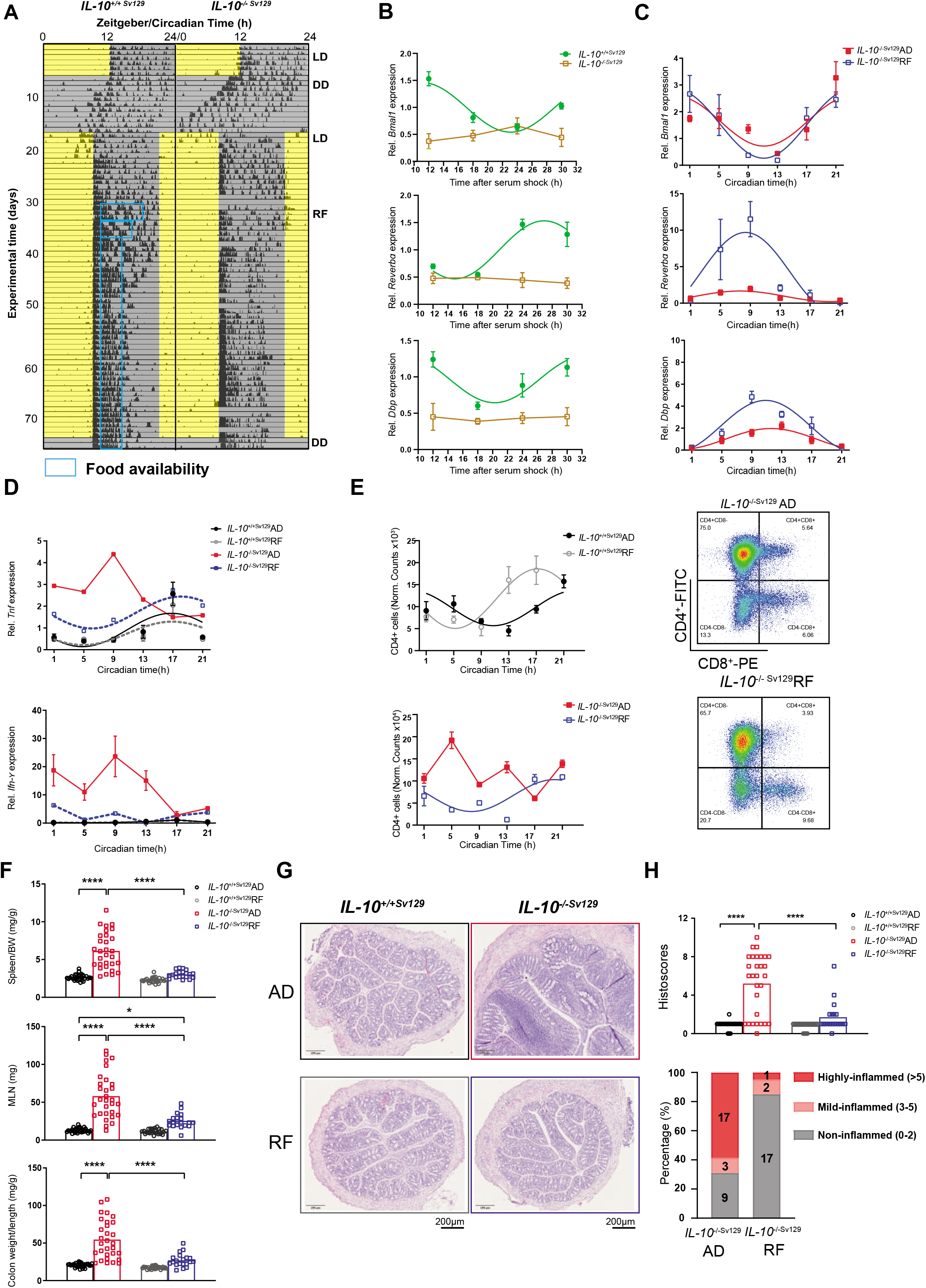
Night time restricted feeding restores the disrupted colonic clock in *IL-10*^-/-Sv129^ mice and ameliorates the colitis. (A) Representative actogram of wild type (*IL-10*^+/+Sv129^) and *IL-10*^-/-Sv129^ mice exposed to light-dark cycle (LD), constant darkness (DD), and night restricted feeding (RF). Tick marks represent running wheel activity; yellow and grey shadings represent light and darkness respectively; blue boxes indicate the time period of food access during RF. (B) Expression of major clock genes *Bmal1*, *Reverba* and *Dbp* in colonic organoids from *IL-10*^-/-Sv129^ mice after synchronization by serum shock. (C) Expression of major clock genes *Bmal1*, *Reverba* and *Dbp* in colonic tissue from *IL-10*^-/-Sv129^ mice under *ad libitum* (AD) and RF conditions. (D) Inflammatory genes expression profile of *Tnf* and *Ifn-y* in colonic tissues from wild type and *IL-10*^-/-Sv129^ mice under AD and RF conditions. (E) Quantification of the amount of CD4+ T cells in the colon lamina propria from wild type and *IL-10*^-/-Sv129^ mice under AD and RF conditions measured by flow cytometry (top) and representative flow cytometric plots for CD4^+^ and CD8^+^ cells (bottom). (F) Organ weights of spleen, mesenteric lymph nodes (MLN) and colon from wild type and *IL-10*^-/-Sv129^ mice in AD and RF conditions. (G) Representative colonic cross section stained with Hematoxylin&eosin (H&E) from wild type and *IL-10*^-/-Sv129^ mice in AD and RF conditions. (H) Histopathological scores of colonic tissues from wild type and *IL-10*^-/-Sv129^ mice in AD and RF conditions (left) and the quantification of individual mice which characterized with different inflammation degrees (right). Tissues with scores between 0-2 are classified as non-inflamed, 3-5 as mildly inflamed and >5 as highly inflamed. Significant rhythms are illustrated with fitted cosine-wave regression using a line (significance: cos-fit p-value ≤ 0.05). Statistics were performed by two-way ANOVA followed with Turkey correction. Data are represented as mean ± SEM. Asterisks indicate significant differences *p<0.05, **p<0.01, ***p<0.001.

Recently we demonstrated an imbalance of the GI immune homeostasis in a genetic mouse model lacking a functional intestinal clock^21^. This prompted us to characterize the molecular intestinal clock in inflamed *IL-10^-/-^* ^Sv129^ mice. Robust circadian rhythms were found in the expression of the essential clock genes *Bmal1* (*Arntl*), *Rev-erbα* (*NR1D1*) and *Dbp* in cultured jejunal and colonic organoids from wild types (*IL-10*^+/+Sv129^). Interestingly, clock gene expression was arrhythmic and suppressed in colonic organoids from inflamed *IL-10-*deficient mice on Sv129 background (*IL-10^-/-^* ^Sv129^) (**Fig. 1B**), indicating a dysfunctional colonic clock. In contrast, clock gene rhythms in jejunal organoids were comparable between genotypes **(Suppl. Fig. 1E)**. These data suggest that circadian dysfunction is restricted to inflamed regions of the intestine. Similar to results obtained from organoids, the rhythmicity of clock gene expression was dramatically altered in colonic tissues from 20 weeks old *IL-10^-/- Sv129^* mice. A genotype difference was identified for *Bmal1* (p=0.0315), and the amplitude of *Rev-Erbα* (p<0.0001) and *Dbp* (p=0.0003) expression were strongly reduced in colonic tissue of *IL-10^-/-^* ^Sv129^ mice compared to wild type littermates, although low amplitude oscillations persist (**Suppl. Fig. 1F, right**). Interestingly, in parallel to disruption of the colonic clock in *IL-10^-/-^* ^Sv129^ mice, complete loss of rhythmicity of lymphocyte recruitment to the lamina propria. CD3^+^CD4^+^ and CD3^+^CD8^+^ T-cell recruitment was observed, whereas immune cell recruitment underwent circadian oscillation in wild types (**Fig. 1E, Suppl. Fig. 1G**). Furthermore, the peak expression of most clock genes was strongly correlated to the colonic histological scores, frequency of immune cells in the lamina propria, including macrophages, neutrophils, CD4^+^ cells and dendritic cells, and the expression of genes involved in barrier function (**Suppl. Fig. 1H-I**). These results indicate a potential causal relationship between colon clock dysfunction in *IL-10*^-/-^ mice and their inflammatory phenotype.

### Night-time restricted feeding restores the colonic clock and ameliorates pathological changes in *IL-10^-/-^* mice

Night-time restricted feeding (RF) is a widely-studied approach in mice to influence the rhythmicity of peripheral clocks^26^. This prompted us to test whether RF in *IL-10*^-/- Sv129^ mice restores the colonic clock and impacts their Crohn’s disease-like phenotype. For this purpose, 14 weeks old *IL-10*^-/- Sv129^ mice were gradually introduced to RF (**Fig. 1A**, **Suppl. Fig. 1A**). Food intake and night-time activity levels during RF were comparable between genotypes (**Suppl. Fig. 1D, E**). Tissues were harvested at the end of the 4-week-RF period during the 2^nd^ day in DD. Indeed, RF improved the circadian amplitude and baseline of clock gene expression in *IL-10*^-/- Sv129^ colonic tissue to similar levels observed in control mice under *ad libitum* (AD) condition (*Rev-erba* p<0.0001, *Dbp* p<0.0001) (**Fig. 1C**, **Suppl. Table 1**). Of note, RF also enhanced the amplitude of clock gene expression in wild types *(Rev-erba* p<0.0001, *Dbp* p=0.0313) (**Suppl. Fig. 1B, left**). Importantly, restoration of the colonic clock in *IL-10^-/- Sv129^* mice exposed to RF conditions was reflected by reduced levels of *Tnf* and *Ifn-y* expression and restored rhythmicity of *Tnf* were observed in *IL-10^-/- Sv129^* mice in contrast to enhanced and arrhythmic expression in AD-fed *IL-10^-/- Sv129^* mice (**Fig. 1D**). Reduced inflammation following RF in *IL-10^-/- Sv129^* mice was additionally reflected by restored rhythmicity in the amount of CD3^+^CD4^+^ cells isolated from colonic lamina propria and an overall reduced amount of CD4 T-lymphocytes (**Fig. 1E, right**). In contrast, CD3^+^CD8^+^ cells recruitment remained arrhythmic (**Suppl. Fig. 1G**). Consistently, the weight of spleen, mesenteric lymph nodes (MLN) and colon density which were severely heavier in AD-fed *IL-10^-/- Sv12^*^9^ mice, were significantly reduced during RF and undistinguishable from controls (**Fig. 1F**). Histological analysis of colon cross sections further revealed more than 58% of *IL-10^-/-Sv12^*^9^ mice under AD condition were identified as highly-inflamed (Histoscore>5), which was dramatically reduced to 5% under RF condition (**Fig. 1G, 1H**). These data demonstrate that RF restores disruption of colon clock function and completely ameliorates the immune phenotype in *IL-10^-/-Sv12^*^9^ mice.

### Loss of microbial rhythmicity during colonic inflammation *in IL-10*^-/-^ mice is restored by RF

Gut microbiota dysbiosis has been associated with the development of IBD in humans and mouse models^19, 20^. Previously we provided the novel link between the intestinal clock, microbiome rhythms and GI homeostasis^21^. This prompted us to investigate circadian rhythmicity in microbiota composition and function in inflamed *IL-10*^-/-^ mice. Indeed, microbiota composition differed significantly between genotypes despite the feeding conditions (**Fig. 2A)** and circadian rhythmicity in community diversity (normalized species richness) was abolished in AD-fed *IL-10*^-/-Sv129^ mice (**Fig. 2B)**. In accordance to results obtained from IBD patients^27, 28^, reduced abundance of *Lachnospiraceae* and *Oscillospiraceae* and increased level of *Erysipelotrichaceae* were observed in *IL-10*^-/-^ mice under AD conditions (**Suppl. Fig. 2A**). The genera *Oscillibacter, Eubacterium, Clostridium* and *Pseudoflavonifractor* are among the taxa which significantly differed in their abundance between genotypes (**Suppl. Fig. 2B left, Suppl. Table. 2**). Moreover, we detected significant correlations between disease markers, such as histological score, inflammatory marker gene expression and zOTUs belonging to *Lachnospiraceae* and *Oscillospiraceae* (**Suppl. Fig. 2B right**), which is consistent with our recent findings showing a correlation between *Lachnospiraceae* and active disease in IBD patients^29^. Similarly, the abundance of both major phyla, Bacteroidota and Firmicutes and highly abundant families, including *Lachnospiraceae* and *Oscillospiraceae* as well as zOTUs, followed circadian oscillation in wild types, whereas these rhythms were abolished in AD-fed *IL-10*^-/-Sv129^ mice (**Fig. 2C, D**). Importantly, RF not only restored abundance differences between genotypes (**Suppl. Fig. 2B**), but also restored microbial rhythmicity in *IL-10*^-/-Sv129^, illustrated in community diversity and on the level of major phyla, the two major families *Lachnospiraceae* and *Oscillospiraceae* (**Fig. 2B, C, E)**. Enhanced microbial rhythmicity was also found among the 237 identified zOTUs visualized by heatmaps **(Fig. 2D, Suppl. Fig. 2C**). More than 50% of the zOTUs which gained rhythmicity during RF in *IL-10*^-/-^ mice belong to the family *Lachnospiraceae* (**Suppl. Table. 2**), which includes bacteria taxa important for SCFA production and secondary bile acids (BAs) conversion and thus plays crucial roles in IBD progression^30^.

**Figure 2.**
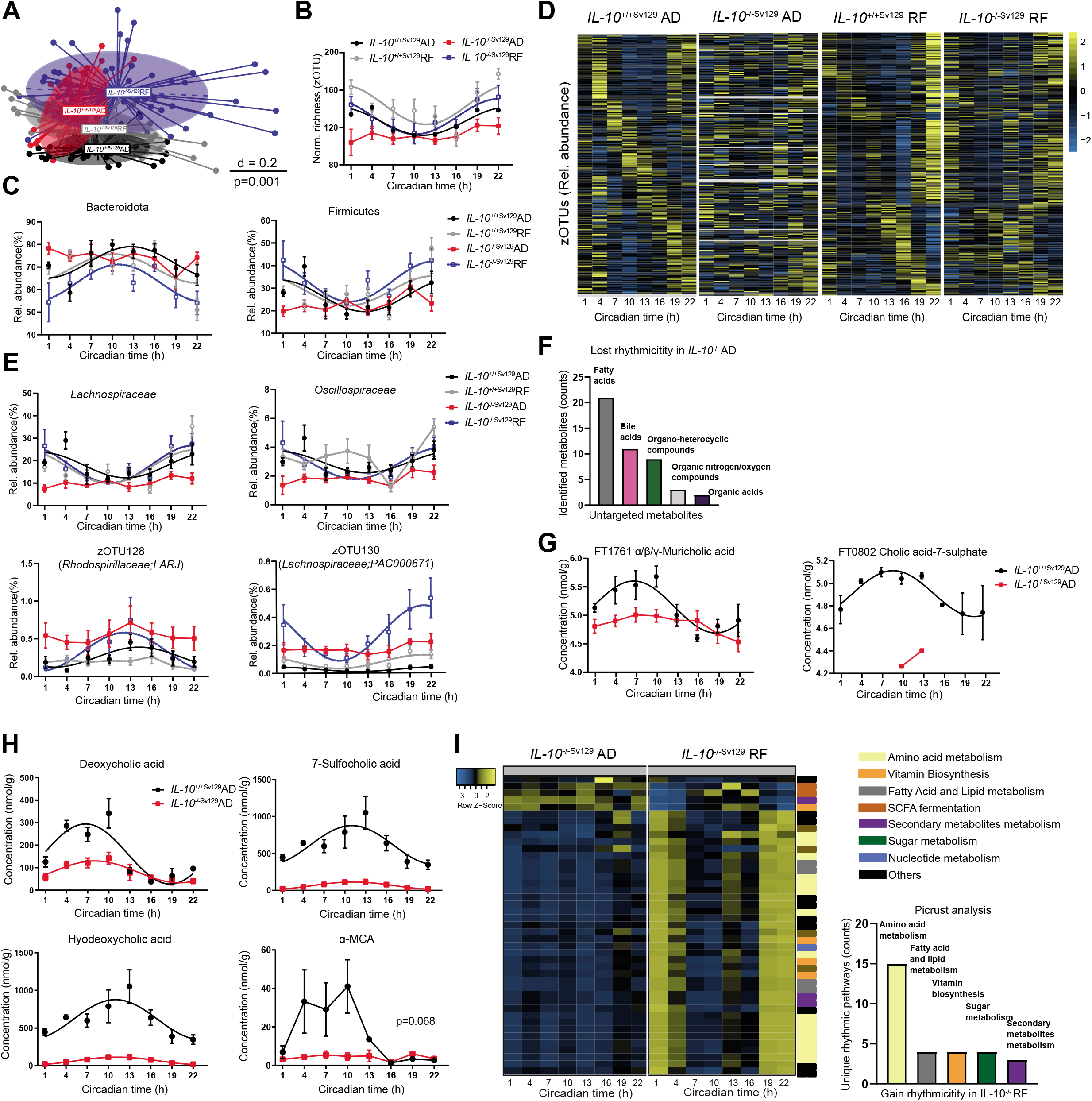
Impaired microbiome rhythms in *IL-10*^-/-SV129^ are restored after RF. (A) Beta-diversity illustrated by MDS plots of fecal microbiota based on generalized UniFrac distances (GUniFrac) in wild type and *IL-10*^-/-Sv129^ mice under AD and RF conditions. Comparison of genotype and feeding effects was performed by PERMANOVA (adj.p=0.001) (B) Circadian profile of normalized richness and (C) relative abundance of two major phyla (*Bacteroidota* and *Firmucutes*) in wild type and *IL-10*^-/-Sv129^ mice under AD and RF conditions. (D) Heatmap depicting the relative abundance of 237 zOTUs (mean relative abundance>0.1%; prevalence > 10%). Data from *IL-10*^-/-Sv129^ mice under AD or RF condition is normalized to the peak of each zOTU and ordered by the peak phase in wild type mice under AD or RF conditions respectively. Yellow and blue indicate high and low abundance respectively. (E) Relative abundance of highly abundant family *Lachnospiraceae* and *Oscillospiraceae* and relative abundance of two representative zOTUs from wild type and *IL-10*^-/-Sv129^ mice under AD and RF conditions. (F) Classification of the top 5 fully annotated super classes of metabolites measure by untargeted metabolomics and (G) representative examples in wild type and *IL-10*^-/-Sv129^ mice under AD condition. (H) Circadian profiles of bile acids (BAs) of wild type and *IL- 10*^-/-Sv129^ mice under AD condition. (I) Heatmap of pathways (calculated by PICRUST 2.0) with restored rhythms *IL-10*^-/-Sv129^ mice under RF (left) and the quantification of the superclass (right). Rhythmicity was identified by JTK_Cycle (Bonferroni adj. p-value ≤ 0.05). Significant rhythms are illustrated with fitted cosine-regression; data points connected by straight lines indicate no significant cosine fit curves (p > 0.05) and thus no rhythmicity. Data are represented as mean ± SEM.

To address the potential physiological relevance of disturbed microbial oscillations in *IL-10*^-/- Sv129^ mice, fecal (un)targeted metabolite analysis was performed. Indeed, the identified metabolites clustered according to the genotype (**Suppl. Fig. 2D**). Moreover, *IL-10*^-/-Sv129^ mice showed loss of rhythmicity in 55 fully annotated metabolites, most of which belong to lipids and lipid-like metabolites, including fatty acids and BAs (**Fig. 2F**) For example, we found loss of rhythmicity and suppression of bile acid, including muricholic acid and 7-sulfocholic acid, as well as the long chain fatty acid oleic acid, linoleic acid (**Fig. 2G, Suppl. Fig. 2E)**. Targeted metabolomics for SCFAs, desaminotyrosine and bile-acids (BAs) further validated the differences in rhythmicity and overall concentrations between genotypes **(Fig. 2H, Suppl. Fig. 2F)**. For example, highly suppressed amplitude rhythms or loss of rhythmicity were observed for primary and secondary BAs in *IL-10*^-/-sv129^ mice, such as deoxycholic acid (DCA) (p<0.0001), hyodeoxycholic acid (p=0.0025), 7-sulfocholic acid (7-sulfo-CA)(p<0.0001) and α-muricholic acid (p=0.0004) (**Fig. 2G**). Most of these secondary BAs are key mediators involved in gut dysbiosis, especially in IBD^31^. Additionally, reduced SCFA levels, such as acetate (p=0.0221), butyrate (p<0.0001) and propionate (p<0.0001) were found in *IL-10*^-/-sv129^ mice **(Suppl. Fig. 2F)**, similar to observation on IBD-patients and related mouse models^32^. The concentration of valeric acid and desaminotyrosin were also suppressed in *IL-10*^-/-Sv129^ mice (**Suppl. Fig. 2F).** Importantly, PICRUSt2 analysis^33^ revealed that restoration of microbial rhythmicity in *IL-10*^-/-Sv129^ mice under RF conditions was indeed reflected by restored rhythmicity of assigned pathways involved in fatty acid synthesis and SCFA fermentation (**Fig 2I**). Altogether, these results indicate that changes in microbial rhythmicity and composition observed in *IL-10*^-/-Sv129^ mice can be restored by RF, and thus microbial rhythmicity might be involved in the beneficial effect of RF on colonic inflammation.

### Germ free intestinal clock-deficient mice develop an increased inflammatory response following transfer of *IL-10*^-/-^-associated microbiota

Recently we demonstrated that a functional intestinal clock is required to maintain GI homeostasis by driving the microbiome, including *Lachnospiraceae* and microbiota derived DCA and 7-sulfo-CA^21^. To differentiate whether a dysfunctional intestinal clock in *IL-10^-/-^*^Sv129^ mice or arrhythmicity of the microbiome directly affect the severity of inflammation, we performed two distinct cecal microbiota transfer experiments. First colonialization of cecal content from intestine-specific clock deficient (*Bmal1*^IEC-/-^) and littermates (*Bmal1*^flox/flox^) donors was performed in order to transfer rhythmic and arrhythmic microbiota into Germ free (GF) *IL-10*^-/-BL^^6^ recipients (**Fig. 3A)**. Surprisingly, the weight of immune organs, such as spleen, MLN and colon did not differ between recipients, nor did the amount of immune cells that infiltrated into the colonic lamina propria (CD4^+^ T-cells, macrophages and neutrophils) (**Fig. 3B**). Consistently, histological staining and scoring, as well as the expression of inflammatory marker gene *Tnf* reveal no pathological differences between both recipients (**Fig. 3C**), indicating that microbial rhythmicity might not directly induce intestinal inflammation. However, when GF *Bmal1*^IEC-/-^ mice received disease-associated microbiota from inflamed *IL- 10*^-/-BL^^6^ mice, colon weights were significantly increased in comparison to control recipients (**Fig. 3D,E**). Similarly, immune cell recruitment into the colonic lamina propria and *Tnf* gene expression was severely elevated, although the histological score obtained from a single colon cross section showed no clear difference between the recipients (**Fig. 3F, G, Suppl. Fig. 3A- C**). Interestingly, we observed beta diversity clustering between circadian time points and recipients (**Fig. 3H**), as well as decreased abundance of *Lachnospiraceae* and increased *Erysipelotrichaceae* in cecal samples in *Bmal1*^IEC-/-^ recipients, similar to results obtained from *IL-10*^-/-Sv129^ mice (**Fig. 3I, Suppl. Fig. 2A**). Accordingly, the concentrations of α-MCA, DCA, HDCA and MDCA in the cecal content were reduced in comparison to control recipients (**Fig. 3J**, **Fig. 2H**). Procrustes analysis (PA) revealed an association between zOTUs with BAs production, in particular the families *Lachnospiraceae* and *Oscillospiraceae* correlated with

**Figure 3.**
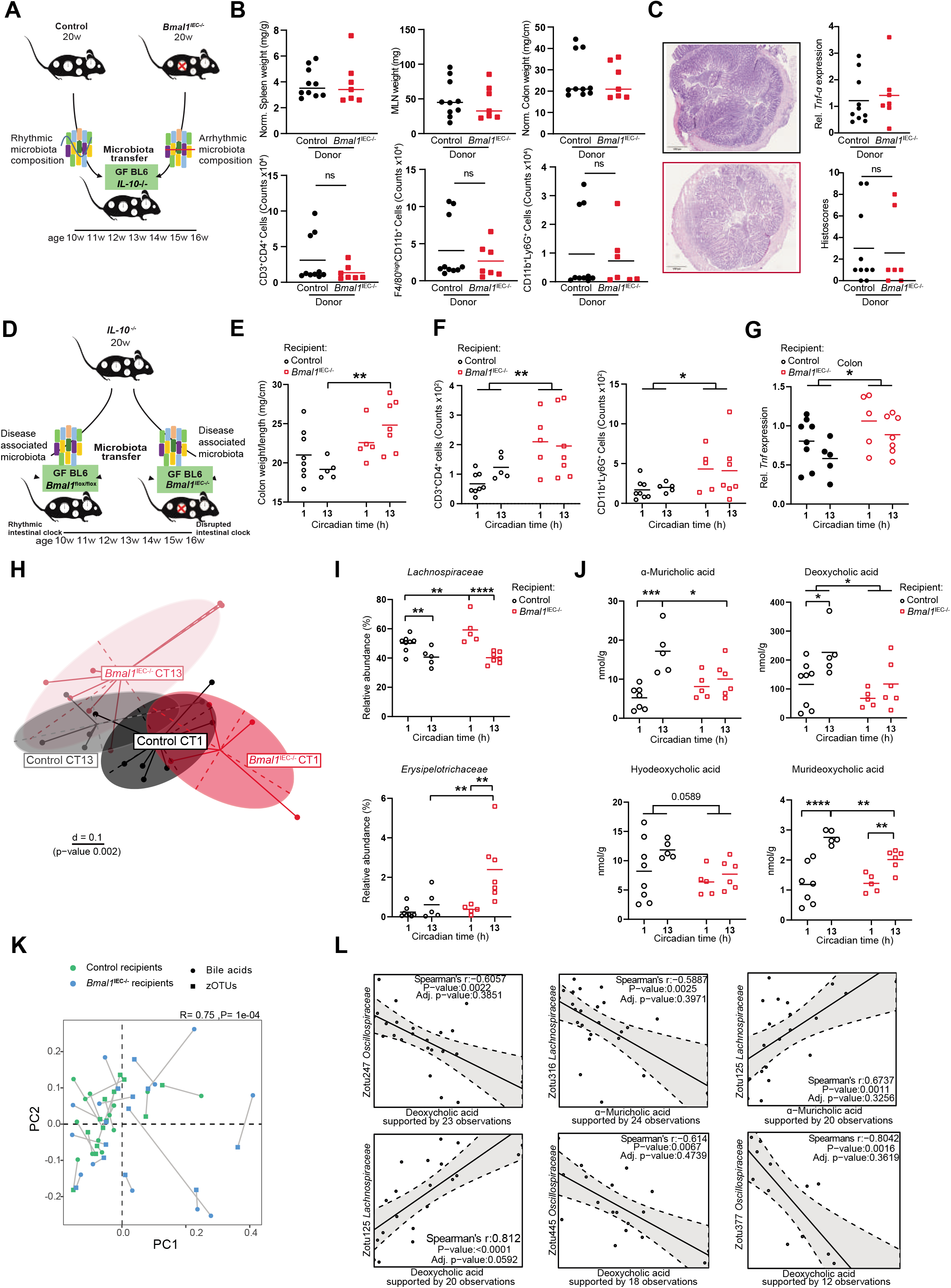
Dysfunction of the host’s intestinal clock promotes microbiota-induced colonic inflammation. (A) Schematic illustration of experimental design. Cecal content from control (rhythmic) and *Bmal1*^IEC-/-^ (arrhythmic) donors were transferred into germ-free *IL-10*^-/-BL^^6^ mice. (B) Organ weights of spleen, MLN and colon (top) and amount of immune cells recruited into colonic lamina propria (bottom) of recipients. (C) Representative H&E stainings of colonic cross section from the recipients (left) and *Tnf* gene expression, as well as histopathological scores (right). (D) Schematic illustration of experiment design. Disease-associated microbiota from *IL-10*^-/-^ donor was transferred to germ-free control (*Bmal1*^flox/flox^) and *Bmal1*^IEC-/-^ recipients. (E) Colon weight and (F) amount of CD4+ T cells and neutrophils recruited into colon lamina propria. (G) *Tnf* gene expression in colon tissues from recipients. (H) Beta-diversity illustrated by MDS plots of cecal microbiota based on generalized UniFrac distances (GUniFrac) in control and *Bmal1*^IEC-/-^ recipients at circadian time (CT) 1 and 13. (I) Relative abundance of families *Lachnosiraceae* and *Erysipelotrichaceae*. (J) Amount of α-muricholic acid (α-MCA), deoxycholic acid (DCA), hyodeoxycholic acid (HDCA) and murideoxycholic acid (MDCA) in cecal content of control and *Bmal1*^IEC-/-^ recipients. (K) Procrustes analysis (PA) of cecal microbiota and bile acid levels. The length of the line is proportional to the divergence between the data from the same mouse. (L) Representative correlation plot between zOTUs and bile acids. Statistics were performed by Mann–Whitney U test and two-way ANOVA followed with Benjamini-Hochberg correction. Asterisks indicate significant differences *p<0.05, **p<0.01, ***p<0.001. Significant rhythms are illustrated with fitted cosine-regression; data points connected by straight lines indicate no significant cosine fit curves (p > 0.05) and thus no rhythmicity. Data are represented as mean ± SEM.

DCA and aMCA concentrations (p=0.0001) (**Fig. 3K, L**). Altogether, these results reflect an early stage of inflammation following colonialization with disease-associated microbiota when recipients lack a functional intestinal clock. Thus, the intestinal clock represents a functional link between the microbiome and GI inflammation.

### Genetic intestinal clock dysfunction alters colonic immune functions

To identify whether intestinal clock dysfunction influences the colonic immune response, we used a genetic mouse model lacking the essential clock gene *Bmal1* in intestinal epithelial cells (IECs) (*Bmal1*^IEC-/-^). Colonic clock dysfunction in IECs was validated by arrhythmic *Bmal1* and *Rev-erba* (*Nr1d1)* expression and a highly reduced amplitude of *Per2* gene expression (**Fig. 4A)**. Furthermore, bulk RNA-sequencing analysis on whole colonic tissue samples obtained during the circadian day revealed that more than 200 genes were differentially expressed (Fold change>1.5, adj. p<0.05) between genotypes (**Fig. 4B**). For example, altered expression was found for genes, such as *Retnlb* (Resistin-like beta) and *Tff*2 (Trefoil factor 2)(**Fig. 4B**) involved in colitis development^34, 35^. Consistently, gene ontology enrichment analysis highlighted pathways relevant for ‘humoral immune response’, ‘defense response to bacterium’ and ‘T-cell apoptotic process’ (**Fig. 4C**). Consistently, clustering according to the genotype was observed by principal component analysis (PCA) (**Fig. 4D**). Moreover, 7803 transcripts underwent circadian rhythmicity in controls, whereas more than 55% of these transcripts were arrhythmic in *Bmal1*^IEC-/-^ mice (**Fig. 4E left, Suppl. Table. 3**). Additionally, the majority of the transcripts which maintained rhythmicity in *Bmal1*^IEC-/-^ mice showed a highly suppressed amplitude (**Fig. 4E, right**). DODR analysis (Pelikan, Herzel et al. 2021) further confirmed that more than 200 genes either lost or changed rhythmicity (114/104 respectively) in *Bmal1*^IEC-/-^ mice and are enriched in pathways including circadian rhythm, inflammatory response and cholesterol homeostasis, similar to results observed between genotype comparison (**Fig. 4C, F**). Particularly, genes such as *Ffar2*, *Abcc2*, *Rorc* and *Hc* which are involved in the inflammatory and immune response^36–39^ lost rhythmicity and in addition significantly differed in their baseline expression between genotypes (**Fig. 4G, Suppl. Table. 3**). Moreover, genes involved in goblet cells function, including *Muc3*, *Tff3*, *Pigr* and *Fcgbp,* lost rhythmicity in *Bmal1*^IEC-/-^ mice (**Suppl. Fig. 4A**)^40^. Genes relevant for cell development and migration, especially immune cells, such as *Casp4*, *Pdk1*, *Cxcr4 and Vcam-1*^41–44^ showed an abnormal circadian phase or loss of rhythmicity upon intestinal-clock disruption (**Suppl. Fig. 4B**). Accordingly, the frequency of major immune cell populations in the colonic lamina propria, including CD4^+^, CD8^+^ T-cells and dendritic cells lost rhythmicity in *Bmal1*^IEC-/-^ mice (**Fig. 4H**), similar to the results obtained from *IL-10*^-/-Sv129^ mice (**Fig. 1E**). Whereas macrophage recruitment remained rhythmic (**Fig. 4H**). Although lack of the intestinal clock in *Bmal1*^IEC-/-^mice does not result in a pathophysiological phenotype (**Suppl. Fig. 4C-E**), colon weight as well as the concentration of complement 3 in stools of *Bmal1*^IEC-/-^ mice was significantly increased (**Fig. 4I**). These findings are consistent with results obtained from *IL-10*^-/-^ mice **(Fig. 1F, Suppl. Fig. 4F**), suggesting that intestinal clock dysfunction might indeed magnify inflammatory processes in *IL-10*^-/-^ mice.

**Figure 4.**
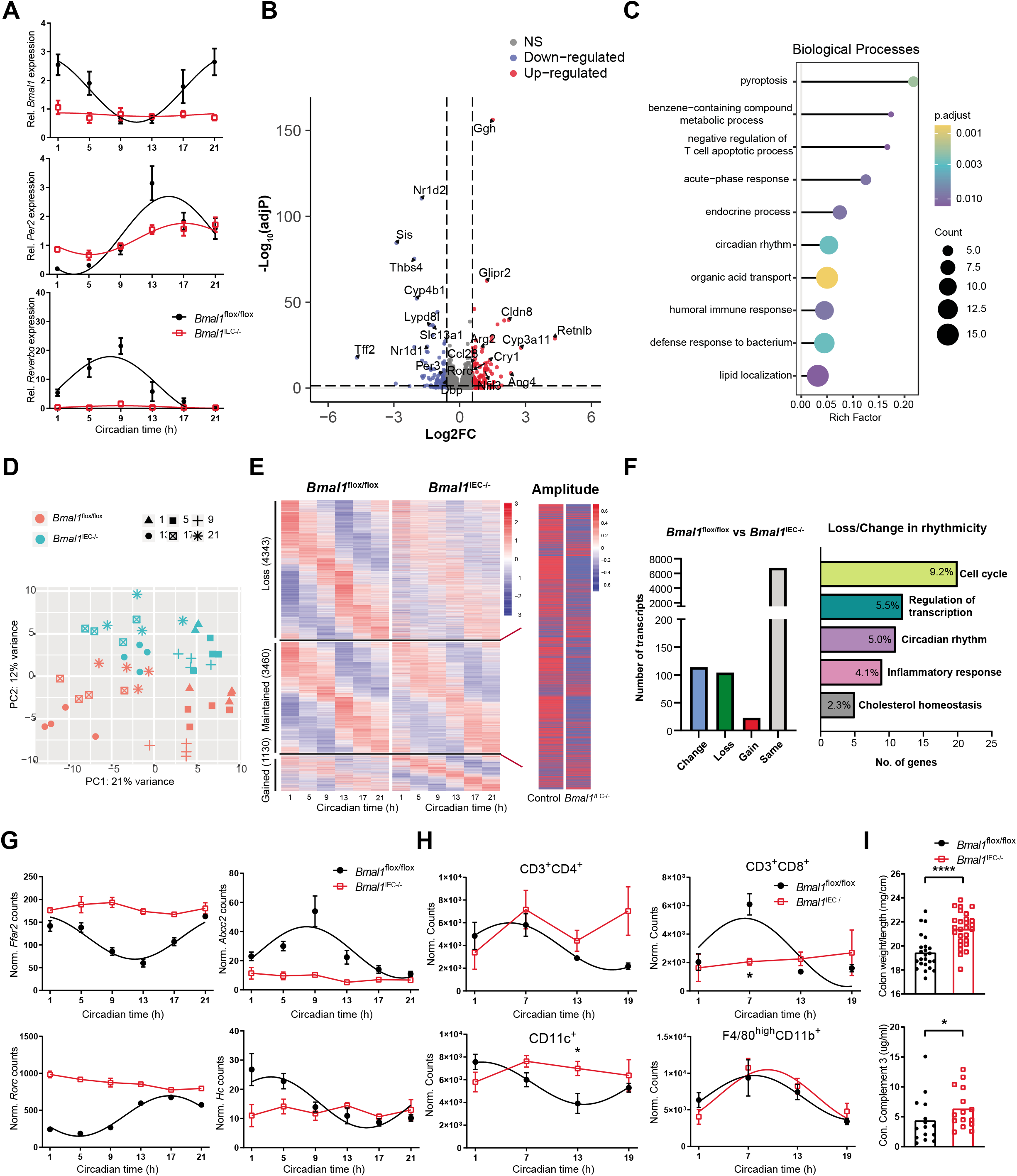
The intestinal clock regulates transcripts involved in intestinal immune responses. (A) Clock genes expression (*Bmal1*, *Per2*, *Reverba)* in colonic epithelial cells from control and *Bmal1*^IEC-/-^ mice. (B) Volcano plot of transcriptional changes between colon tissues from control and *Bmal1*^IEC-/-^ mice. Upregulated (adj p < 0.05, FC > 1.5) genes are indicated in red, while downregulated (adj p < 0.05, FC > 1.5) genes are marked in blue. (C) Gene Ontology enrichment analysis of differentially expressed genes. (D) Principal Component Analysis of 46 colon samples obtained from control (red) and *Bmal1*^IEC-/-^ (blue) mice every 4h throughout the circadian day in constant darkness. Different shapes indicate different circadian time (CT) points. (E) Heatmap of transcripts which lost, maintained and gained rhythmicity in colon tissues from *Bmal1*^IEC-/-^ mice (left) and amplitude comparison of the transcripts which maintained rhythmicity (right) (JTK_cycle adj.p <0.05 as rhythmic). Transcripts are ordered by the peak phase of control groups. (F) Amount of transcripts calculated by compareRhythms (left) and GO analysis (top 5, right) which loss and change rhythmicity. (G) Circadian profile of *Ffar2*, *Abcc2*, *Rorc* and *Hc*. (H) Amount of major subtypes of immune cells in the colon lamina propria of control and *Bmal1*^IEC-/-^ mice. (I) Colon weight (top) and complement 3 level in feces of control and *Bmal1*^IEC-/-^ mice. Statistics were performed by Mann–Whitney U test. Asterisks indicate significant differences *p<0.05, **p<0.01, ***p<0.001.

### *Bmal1*^IEC-/-^ mice are more susceptible to acute and chronic colon inflammation

In order to directly test the cause-effect relationship between intestinal clock dysfunction and GI inflammation, *Bmal1*^IEC-/-^mice and controls were released into DD and received drinking water supplemented with 2% dextran sulfate sodium (DSS) (**Fig. 5A**), a well-studied chemical to induce tissue damage followed by colitis^45^. 5 days of DSS treatment caused a significant reduction in body weight and an increased disease activity index (DAI) ^46^ (**Fig. 5B,C**), as well as shortened colon length (**Fig. 5D**) compared to DSS-treated controls. Accordingly, the expression of inflammatory markers, such as *Tnf* as well as neutrophil and macrophage recruitment to the colonic lamina propria was severely enhanced following DSS treatment in *Bmal1*^IEC-/-^ mice (**Fig. 5E-F**), suggesting an increased inflammatory response. Consistently, the histopathological evaluation in colon tissues significantly differed between genotypes after DSS treatment, which was reflected by an increased histological scores in *Bmal1*^IEC-/-^mice **(Fig. 5G-H**). Additionally, intestinal clock dysfunction caused loss of goblet cells following DSS treatment (**Fig. 5I**), which was previously identified as a critical factor during DSS-induced colitis^46^. These results demonstrate a higher sensitivity to DSS-induced colitis in mice lacking a functional intestinal clock.

**Fig. 5.**
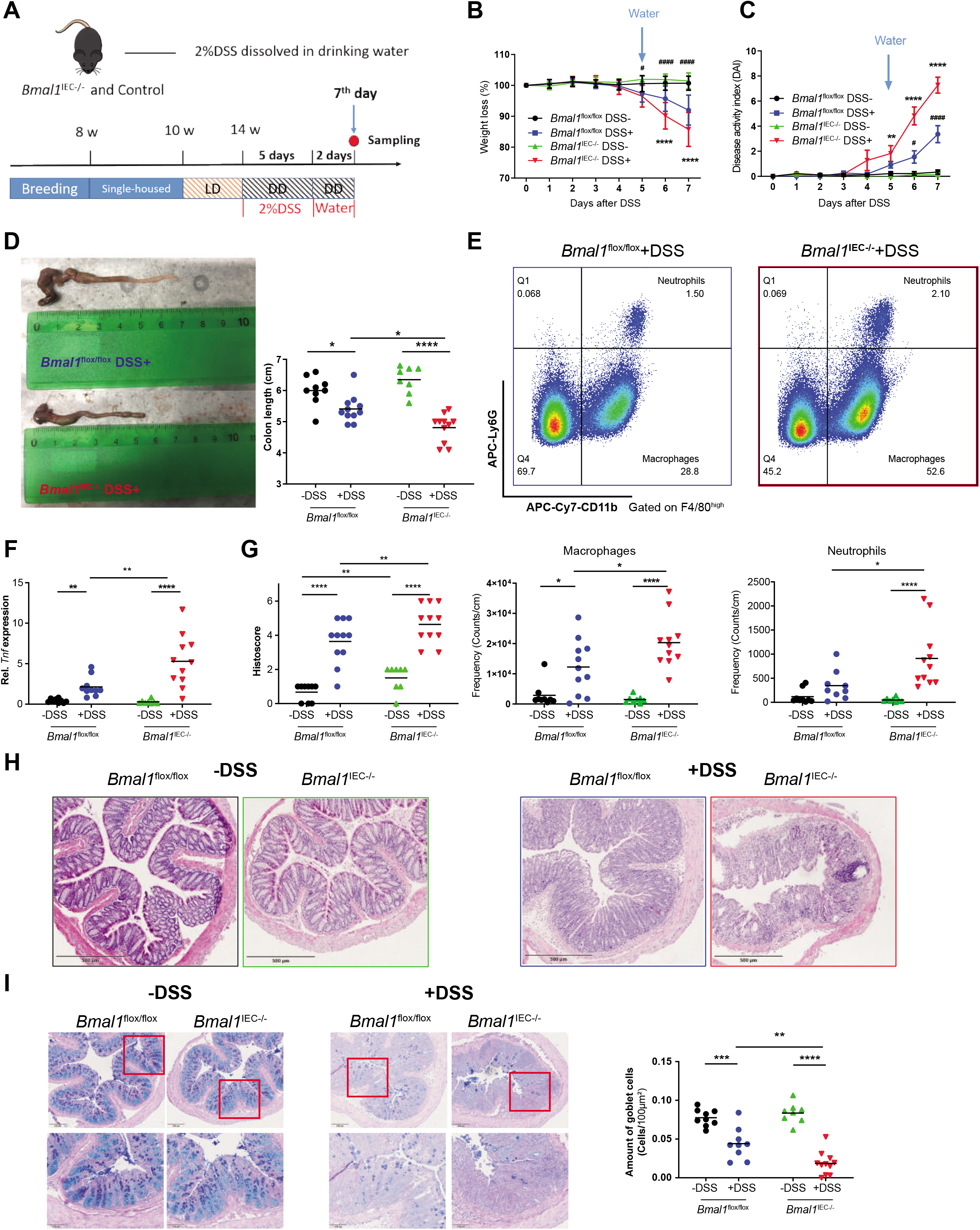
*Bmal1*^IEC-/-^ mice are more susceptible to DSS-induced colitis. (A) Schematic illustration of the DSS treatment procedure. (B) Body weight change and (C) disease activity index of control and *Bmal1*^IEC-/-^ during DSS/water administration. (D) Representative picture of colon (left) and colon length measurement (right). (E) Gating strategy for macrophages and neutrophils in colon lamina propria (top) and quantification (bottom). (F) *Tnf* expression of colon tissues. (G) Histopathological scores, (H) representative H&E staining (I) and PAS-AB staining of colon sections from control and *Bmal1*^IEC-/-^ with/without DSS treatment and quantification of amount of goblet cells. Quantification was based on a minimum of 20 crypts for each section. Statistics were performed by two-way ANOVA followed with Benjamini-Hochberg correction. Asterisks indicate significant differences between *Bmal1*^flox/flox^ and *Bmal1*^IEC-/-^ mice after DSS treatment *p<0.05, **p<0.01, ***p<0.001. Octothorpes indicate significant differences between *Bmal1*^flox/flox^ mice with and without DSS treatment #p<0.05, ##p<0.01, ###p<0.001.

In order to genetically assess the impact of the intestinal clock on IBD development, we determined IBD pathology in a newly generated genetic mouse model prone to develop chronic colitis combined with a dysfunctional intestinal clock (*Bmal1*^IEC-/-^ and *IL-10*^-/-BL^^6^). IL-10- deficiency on BL6 background causes less severe colitis symptoms than on SV129 background ^25^, which is likely a results of less severe alterations in clock gene expression (**Fig. 6A**, **Suppl. Fig. 1B, Suppl. Fig. 5F**). Accordingly, the additional loss of the intestinal clock dramatically reduced the survival of *Bmal1*^IEC-/-^x*IL-10*^-/-BL^^6^ mice (59%) compared to *IL-10*^-/-BL^^6^ mice (81.8%) (**Fig. 6B**). In particular, all *Bmal1*^IEC-/-^ x *IL-10*^-/-BL^^6^ mice needed to be euthanized before the end of the experiment due to their dramatic disease burden, while around 30% of *IL-10*^-/-BL^^6^ mice survived. Consequently, severer tissue inflammation was observed in *Bmal1*^IEC-/-^x*IL-10*^-/-BL^^6^ mice, which was reflected by an increased spleen, colon and MLNs weight and a higher tissue inflammation (**Fig. 6C, D**). Similar to our previous results obtained from *Bmal1*^IEC-/-^ mice^21^, circadian rhythmicity in community diversity (species richness) and on the level of the major phyla was dramatically disrupted in *Bmal1*^IEC-/-^ x *IL-10*^-/-BL^^6^ mice, whereas *IL-10*^-/-BL^^6^ mice showed slightly altered circadian oscillations of richness and the major phyla (**Suppl. Fig. 5A- C)**. This genotype effect was even more pronounced at the level of bacterial taxa. A heatmap showing peak relative abundances of zOTUs and its quantification confirmed the disruption of microbial rhythmicity in *Bmal1*^IEC-/-^x*IL-10*^-/-BL^^6^ mice, whereas intermediate phenotypes were noted in *IL-10*^-/-BL^^6^ mice (Suppl. **Fig. 5D, E**). Altogether these data demonstrate that lack of the intestinal clock promotes the sensitivity to acute and chronic colon inflammation.

**Figure 6.**
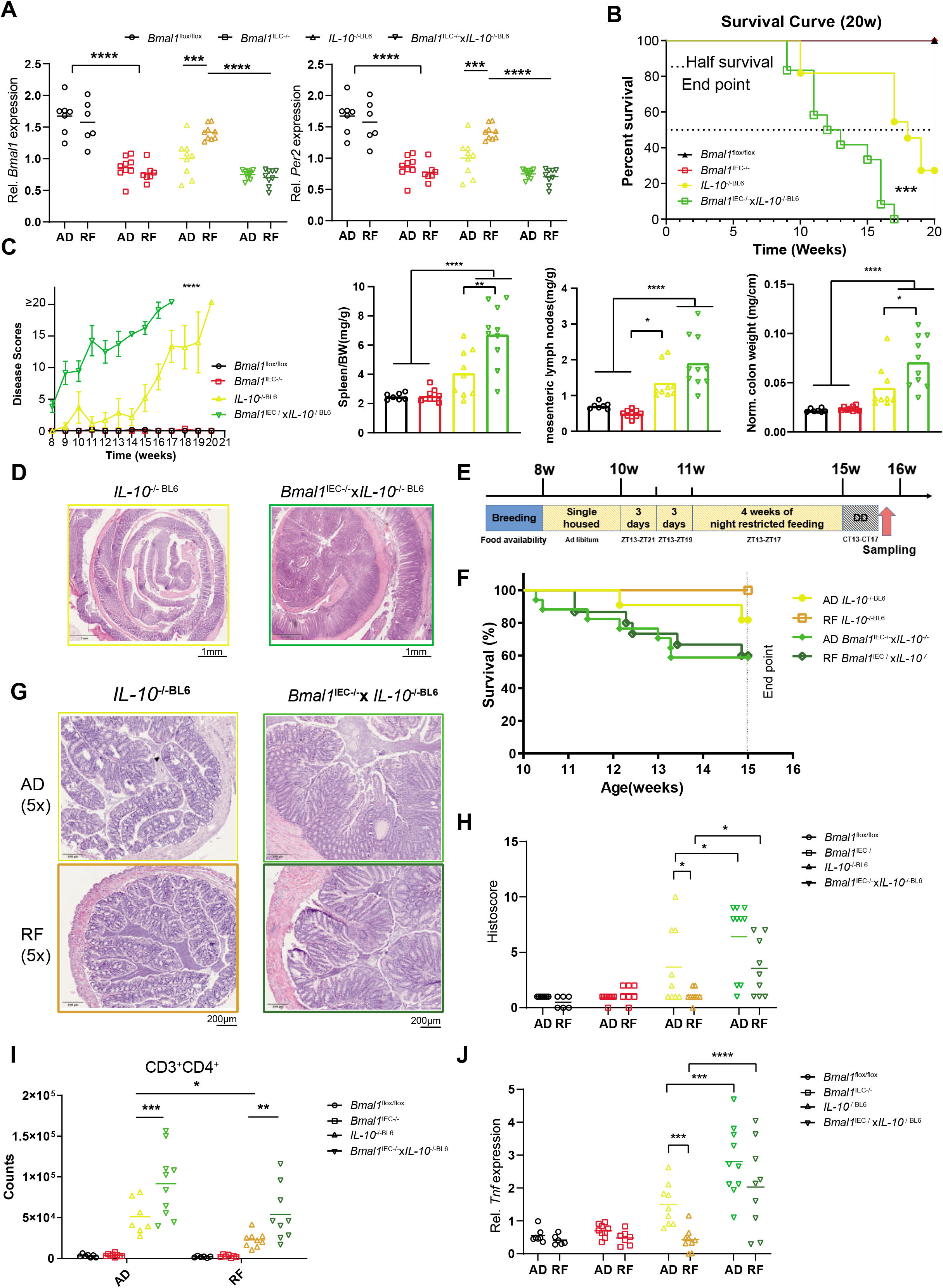
Night time restricted feeding ameliorate the IBD-like symptoms in mice by targeting the intestinal clock. (A) Colon clock genes expression and of control, *Bmal1*^IEC-/-^, *IL-10*^-/-BL^^6^ and *Bmal1*^IEC-/-^x*IL-10*^-/-BL^^6^ mice under AD and RF conditions. (B) Survival analysis of control, *Bmal1*^IEC-/-^, *IL-10*^-/-BL^^6^ and *Bmal1*^IEC-/-^x*IL-10*^-/-BL^^6^ mice under AD condition. (C) Disease scores and organ weights of spleen, MLN and colon. (D) Representative H&E stainings of colon swiss rolls from *IL-10*^-/-BL^^6^ and *Bmal1*^IEC-/-^x*IL-10*^-/-BL^^6^ mice under AD condition. (E) Schematic illustration of experimental RF design. (F) Survival analysis of *IL-10*^-/-BL^^6^ and *Bmal1*^IEC-/-^x*IL-10*^-/-BL^^6^ mice under AD and RF condition. (G) Representative H&E stainings of colon section from *IL-10*^-/-BL^^6^ and *Bmal1*^IEC-/-^x*IL-10*^-/-BL^^6^ mice under AD and RF conditions. (H) Histopathological scores and (I) frequency of CD3+CD4 cells in colon lamina propria and (J) *Tnf* expression. Significances were calculated by one-way ANOVA, two-way ANOVA following Benjamini-Hochberg correction, and three- way ANOVA following Benjamini, Krieger and Yekutieli correction. * p ≤ 0.05, ** p ≤ 0.01, *** p ≤ 0.001, **** p ≤ 0.0001. Data are represented as mean ± SEM.

### Restricted feeding requires a functional intestinal clock to ameliorate colitis symptoms

To evaluate whether the beneficial effect of RF on the severity of GI inflammation observed in *IL-10*^-/-Sv129^ mice is caused directly by restoration of gut clock function, *Bmal1*^IEC-/-^x*IL-10*^-/-BL^^6^ mice, *IL-10*^-/-BL^^6^ and controls were exposed to RF conditions (**Fig. 6E**). We validated that following exposure to RF, clock gene expression at CT13 was recovered similar to control levels in *IL-10*^-/-BL^^6^ mice, whereas the expression was not restored in *Bmal1*^IEC-/-^ mice and *Bmal1*^IEC-/-^x*IL-10*^-/-BL^^6^ mice, due to lack of the intestinal *Bmal1* (**Fig. 6A, Suppl. Fig. 5F**). In line with our hypothesis, RF failed to improve the survival of *IL-10*^-/-BL^^6^ mice when the intestinal clock was genetically dysfunctional (60%), whereas survival was significantly improved by RF in *IL-10*^-/-BL^^6^ mice (100%) (**Fig. 6F**). Consistently, the pathohistological evaluation (**Fig. 6G, H**), CD4 T-cell recruitment to the lamina propria and colonic *Tnf* expression were indistinguishable between AD and RF conditions in *Bmal1*^IEC-/-^x*IL-10*^-/-BL^^6^ mice (**Fig. 6I, J**). Interestingly, RF in *Bmal1*^IEC-/-^ x *IL-10*^-/-BL^^6^ mice enhanced species richness to control levels, whereas microbiota rhythmicity remained arrhythmic (**Suppl. Fig. 5G**), indicating that loss of microbial rhythms might be involved in the disease phenotype rather than the overall amount of species. Indeed, we identified 40 zOTUs which gained rhythmicity during RF in *IL-10*^-/-BL^^6^ mice, but remained arrhythmic in *Bmal1*^IEC-/-^ x *IL-10*^-/-BL^^6^ mice. The majority of these zOTUs belong to the family *Lachnospiraceae, Muribaculaceae* and *Oscillospiraceae* (**Suppl. Fig. 5H, Suppl. Table. 4**), which were previously identified as intestinal clock-controlled^21^.

In summary, these results clearly demonstrate that intestinal clock dysfunction significantly contributes to the colitis severity and that RF improved GI inflammation by targeting intestinal clock functions. Moreover, we provide the first evidence that the intestinal clock gates the inflammatory response and can directly be targeted by RF to mediate the severity and progression of IBD symptoms.

## Discussion

Alterations in clock genes were found in colonic biopsy samples from IBD patients, especially patients with ulcerative colitis^22, 23^. In this study on *IL-10^-/-^* mice, a previously-described IBD-relevant mouse model for colitis^25^, we provide evidence that a disrupted colonic circadian clock is a cause rather than the consequence of GI inflammation in IBD. DSS treatment severely enhanced chemically induced colitis in mice lacking a functional intestinal circadian clock. This is in accordance with previous literature which showed that environmental circadian disruption in form of shift work as well as system-wide genetic clock dysfunction, e.g. in *Per1/2^-/-^*, *Rev- erbα*^-/-^ and *Bmal1*^-/-^ mice promotes the severity of DSS-induced colitis^47–49^. Similarly, recent literature indicated disruption of diurnal intestinal rhythmicity contributes to the DSS-induced colitis^50^. However, we demonstrate for the first time that specifically loss of the circadian intestinal clock promotes the severity and prevalence of DSS-induced colitis symptoms in mice. The cause-effect relationship was further validated in a novel genetic mouse model for colitis which additionally lack a functional intestinal clock. These *Bmal1*^IEC-/-^x*IL-10*^-/-BL^^6^ mice develop a severely enhanced immune phenotype and dramatically reduced survival.

RNAseq analysis on mice with genetic dysfunction of the intestinal clock indicated potential mechanisms how a disrupted colon clock in e.g. *IL-10^-/-^* mice might affect GI inflammation. A majority of intestinal-clock controlled genes were found to be involved in local immune functions and epithelial-microbe interactions. The intestinal clock regulates important inflammatory marker genes, such as *Tnf*, which are arrhythmic and enhanced in *IL-10^-/-^* mice and have previously been linked to the development of IBD^51^. Intestinal clock-controlled genes are also involved in immune cell recruitment, such as *Ffar2,* which plays an important role in intestinal homeostasis by regulating immune cell abundance in colonic lymphoid tissues and the secretion of mucus-associated proteins and antimicrobial peptides^36^ as well as *Abcc2,* which actively regulates mucosal inflammation^37^. Both genes are differentially expressed and lost rhythmicity in mice lacking an intestinal clock. Moreover, *Rorc* which encodes the IBD risk factor RORγ^38^, was elevated and arrhythmic in mice lacking intestinal clock. These results further suggest the importance of the intestinal clock in modulating IBD-like phenotype. In addition, the expression of the pro-migratory genes *Vcam-1* and *Cxcr4* undergoes diurnal oscillation in multiple lymphoid and non-lymphoid tissues^52, 53^. Here we further extended these observations to the colon, that the circadian expression of these two genes require a functional intestinal clock. Furthermore, *Cxcr4* mediates the diurnal oscillations in T cell distribution and responses^52^ and *Vcam-1* is involved in the rhythmic pattern of leukocytes migration^53^. Therefore, arrhythmicity of these genes likely contributes to the immune cell recruitment to colon lamina propria and thus impact the local immune response. Indeed, here we provide the first evidence for a functional role of the intestinal clock in regulating rhythmic immune and inflammatory processes including immune cell recruitment and microbiota composition, which are both key elements in IBD development or progression^51^. For example, a dysfunctional colon clock in *IL- 10*^-/-^ and *Bmal1*^IEC-/-^ mice caused loss of rhythmicity in the recruitment of T cells into the colonic lamina propria. Similarly, rhythmic leukocyte trafficking into the bloodstream and tissue infiltration to lymph nodes, spleen and bone marrow has previously been described for T cells and monocytes and depends on cell- and tissue-specific circadian clocks^54^. The physiological importance of diurnal immune cell recruitment was demonstrated for T-cells recruitment in lymphoid organs which improved the immune response to antigens and bacterial infection at night^52^. Moreover, clustering specifically of CD4+ cells in the intestine induced colitis in an adoptive transfer mouse model^55^. Consequently, arrhythmic leukocyte recruitment to the colon might have augmented the inflammatory response in mice with a disrupted colon circadian clock.

Microbiota composition and function has frequently been associated to the development of GI inflammation^56^ and loss of rhythmicity of the microbiome was linked to the development of obesity and Type 2 diabetes^57^. Interestingly, microbiome rhythmicity was disrupted in *IL-10*^-/- Sv129^ mice. This was likely caused by intestinal clock dysfunction in these mice, because the intestinal clock was recently identified as a major driver of microbiota rhythmicity^21^. Similar to previous results obtained from *Bmal1*^IEC-/-^ mice^21^, intestinal clock-controlled taxa involved in SCFA fermentation and secondary bile acid formation, such as *Lachnospiraceae* and *Oscillospiraceae*, were arrhythmic in *IL-10*^-/-Sv129^ mice. Arrhythmicity of these specific taxa in *IL-10*^-/-Sv129^ mice likely caused the reduced amplitudes or loss of rhythmicity in the levels of specific BAs, such as DCA, 7-sulfo CA, HDCA and α-MCA, which have frequently been linked to intestinal inflammation^58, 59^. Notably, reduced levels of secondary BAs, e.g. DCAs and HDCAs identified in *Bmal1^IEC-/-^* mice receiving disease-associated microbiota, were also found in in patients with UC and supplementation particularly with DCA ameliorates experimental colitis in mice^60^. Thus arrhythmic microbial metabolite production due to loss of rhythmic microbiota composition in *IL-10^-/-^* mice might promote their inflammatory immune response. Indeed, our recent transfer experiments provided evidence that arrhythmic microbiota and derived products can alter GI immune homeostasis^21^. However, transfer of arrhythmic microbiota from *Bmal1^IEC-/-^* did not increase the severity of inflammation in germ-free *IL-10^-/-^* recipients, suggesting that intestinal clock dysfunction rather than loss of microbial rhythmicity induce colonic inflammation. Indeed, transfer of IBD-associated microbiota provide direct evidence for the physiological relevance of the intestinal circadian clock for the inflammatory response. Depending on the recipient genotype, disease-associated microbiota caused an inflammatory response, which reflects the one observed in *IL-10*^-/-Sv129^ mice. Enhanced CD4^+^ T-cells infiltration and elevated *Tnf* expression was only found in colonic tissues of *Bmal1^IEC-/-^* recipients. Moreover, similar to *IL-10*^-/-Sv129^ mice, we found altered abundance of zOTUs belonging to *Lachnospiraceae* and *Erysipelotrichaceae*, and IBD-associated microbiota-derived products, such as α-MCA and DCA, were suppressed and their time difference was lost in *Bmal1^IEC-/-^* recipients. These results reflect the immune response observed in the *IL-10*^-/-^ donors and thus indicate an activated, early inflammatory response although in colon cross sections no pathohistological tissue changes could be observed between genotypes. This might be due to the patchy colonic inflammation associated with the Crohn’s disease-like phenotype reported in *IL-10*^-/-^ mice^61^. Altogether, these results highlight the relevance of intestinal clock function for GI inflammatory processes involved in microbiota-induced IBD development. Interestingly, a functional intestinal clock in control recipients was capable of restoring the circadian time differences of zOTUs. These rhythmic hosts showed reduced immune cell recruitment into lamina propria and inflammatory marker gene expression and thus indicates that restoration of the intestinal clock might reduce GI inflammation.

In order to determine whether intestinal clock function can be targeted to influence the immune response to GI inflammation, *IL-10^-/-^* mice were exposed to night time restricted feeding (RF). RF in mice kept in different light conditions was capable to influence the rhythmicity of major clock genes in intestinal tissues independent of the central clock ^62, 63^. Here we demonstrate that the colonic circadian clock can indeed be restored by RF in a diseased-mouse model (*IL-10^-/-^*) with disrupted colonic clock function. Targeting the colon clock by RF further led to restoration of rhythmic colon clock functions, such as CD4^+^ T-cell recruitment into the colonic lamina propria and microbiome oscillations. In confirmation with our hypothesis RF ameliorated the IBD-like colitis phenotype in *IL-10^-/-^* mice and substantially enhanced their survival. This is in accordance with results obtained in mice showing that RF protects from DSS-induced colitis by affecting intestinal functions^64, 65^. Similarly, RF in a mouse model for arthritis influences leukocytes responsiveness and improves systemic inflammation^66^. In humans, meal timing was tested as treatment for a variety of diseases including diabetes, cancer and tissue injuries^67^. Although inconsistent meal times could be correlated to IBD symptoms^68^, to our knowledge no clear evidence has been generated regarding the beneficial effect of RF on IBD patients. Moreover, the mechanism of RF on disease outcome remains unclear. Although our results indicate that RF might prevent or reduce inflammation by activating the intestinal clock, this is particularly difficult to prove due to the circadian system-wide effects of RF. Namely, RF was found to induce phase shifts in clock gene expression in the liver, kidney, pancreas, WAT and intestinal tissue^69, 70^ and to affect gene expression involved in inflammatory signalling such as NF-kB, TLR or IL-17 in many tissues including the intestine^71^. Here, we provide the first direct mechanistic evidence that the effect of RF on the severity of IBD is gated by the intestinal clock, because RF failed to reduce the inflammatory phenotype (immune cells infiltration, inflammatory marker gene expression, histopathological scores) due to genetic dysfunction of the intestinal clock in *Bmal1^IEC-/-^*x*IL-10^-/-^* mice. Our results are in accordance with a report indicating *Bmal1* gene expression in colonic/jejunal IECs could not be restored by RF^72^.

Notably, we observed an improved richness following RF in *Bmal1^IEC-/-^*x*IL-10^-/-^* mice (**Suppl. Fig. 5K**), which has been associated with improved IBD conditions^73^. Thus, we cannot exclude that other factors, such as the microbiome^17, 74^ and hormones^75^, known to respond to RF independent of the intestinal clock, might additionally contribute to the improved disease outcome. Nevertheless, no significant improvements in tissue pathology or the inflammatory response nor enhanced overall survival following RF was observed in *Bmal1^IEC-/-^*x*IL-10^-/-^* mice. Thus, these results demonstrate that the intestinal clock represents a target for treatment of gastrointestinal inflammation.

Taken together, our results demonstrate that a functional intestinal clock is essential to maintain GI homeostasis and represents a major player in IBD progression. In addition, our findings suggest that enhancing intestinal clock function by meal timing might become the next step to develop novel strategies in circadian therapies of IBD and potentially other metabolic diseases in humans.

**ADDITIONAL INFORMATION (CONTAINING SUPPLEMENTARY INFORMATION LINE (IF ANY) AND CORRESPONDING AUTHOR LINE)**

## ACKNOWLEDGEMENT

The Technical University of Munich provided funding for the ZIEL Institute for Food & Health, animal facility support, technical assistance and support for 16S rRNA gene amplicon sequencing. Moreover, SK was supported by the German Research Foundation (DFG, project KI 19581) and the European Crohńs and Colitis Organisation (ECCO, grant 5280024). SK and DH received funding by the Deutsche Forschungsgemeinschaft (DFG, German Research Foundation) – Project number 395357507 – SFB 1371). We are thankful for Kinga Balázs providing assistance with RNA sequencing analysis and Elizaveta Gorbunova providing assistance with animal experiments.

## AUTHOR CONTRIBUTIONS

SK conceived and coordinated the project. YN performed mouse experiments and fecal samples collection. YN sampled tissue, performed gene expression analysis and FACS analysis. YN analyzed 16S rRNA gene sequencing data and conducted bioinformatics. BA performed the transfer experiments and predicted microbiota functionality analysis. MH, CM and KK performed untargeted and targeted metabolomics and data analysis. MB generated preliminary Flow cytometry data. SK analysed activity behavior and supervised the work and data analysis. SK and DH secured funding. YN and SK wrote the manuscript. All authors reviewed and revised the manuscript.

## MATERIALS & CORRESPONDENCE

Correspondence to Dr. Silke Kiessling, Chair of Nutrition and Immunology, Technical University of Munich, Gregor-Mendel-Str. 2, 85354 Freising, Germany.

## METHODS

### Ethics Statement

Experiments were conducted at Technical University of Munich in accordance with Bavarian Animal Care and Use Committee (TVA ROB-55.2Vet-2532.Vet_02-18-14).

### Mouse models

#### *Bmal1*^IEC -/-^ and *Bmal1*^flox/flox^ mouse generation

Male intestinal epithelial cell-specific *Bmal1* knock-out *(Bmal1fl/fl* x Villin CRE/wt; referred to as *Bmal1*^IEC-/^*^-^*) mice and their control littermates (*Bmal1fl/fl* x Villin wt/wt ; referred to as *Bmal1*^flox/flox^) on a genetic C57BL/6J background were generated as previously described^76^. Breeding was performed by crossing *Bmal1fl/fl* x Villin CRE/wt with *Bmal1fl/fl* x Villin wt/wt. Mice were kept in LD 12:12 cycles (300 lux), with lights turned on at 5am (*Zeitgeber* time (ZT0)) to 5pm (ZT12)) unless stated otherwise. At the age of 8 weeks, mice were single housed with running wheel at 22 ± 1 °C. Mice have ad libitum access to chow diet (V1124-300, Ssniff Diets, Soest, Germany) and water under specific-pathogen free (SPF) conditions according the FELASA recommendation unless stated otherwise.

#### *Bmal1*^IEC -/-^x*IL-10*^-/-BL^^6^ mouse generation and *IL-10*^-/-Sv129^ mouse

Male intestinal epithelial cell-specific *Bmal1* and interleukin-10 double knock-out mice (*Bmal1*^IEC -/-^x*IL-10*^-/-BL^^6^) were generated by crossing *Bmal1*^flox/flox^x*IL-10*^+/-BL^^6^ with *Bmal1*^IEC-/-^ x*IL-10*^+/-BL^^6^ under SPF conditions and bred for several generations internally to harmonize the intestinal microbiota. Interleukin-10 knock-out mice under BL6 and Sv129 background were initially provided by The Jackson Laboratory and bred internally.

### Behavior analysis

Handling and activity measurements during experiments were performed as described^77^. Wheel-running activity was analyzed using ClockLab software (Actimetrics). The last 10-14 days of each condition were used to determine the period (tau, calculated using a X^2^ periodogram and confirmed by fitting a line to the onsets of activity), the duration of the active period (alpha), the amount of activity and the subjective day/night activity ratio (where the subjective day under DD conditions is the inactive period between the offset of activity and the onset of activity and the subjective night is the active period between the onset of activity and the offset of activity). Average daily food intake was measured in the second week of different light conditions under *ad libitum* condition. For restricted feeding condition, average daily food intake was measured manually over 3 consecutive days in the last week of restricted feeding period.

#### Light-dark (LD) and constant darkness (DD) conditions

Male *IL-10*^-/-Sv129^ mice and wild type littermates were single-housed at 8 weeks old under LD cycle for 2 weeks (age 8-10 weeks), switched to a DD cycle for 2 more weeks (age 10-12 weeks) and subsequently back to LD.

#### Night Time Restricted feeding

14 weeks old *IL-10*^-/-Sv129^ mice and wild type littermates were gradually introduced to the restricted feeding (RF) protocol starting with 8h food availability (ZT13-ZT21) for the first 3 days, followed by 6h food availability between (ZT13-ZT19) for the following 3 days and finally 4h food availability (ZT13-ZT17) for four weeks. The same RF regimen also applied to *Bmal1*^flox/flox^, *Bmal1*^IEC-/-^, *IL-10*^-/-BL^^6^ and *IL-10*^-/-BL^^6^ x *Bmal1*^IEC-/-^ mice, with an earlier start point at age of 10 weeks. Tissues were harvested at the end of the 4-week-RF period, on the 2nd day in constant darkness.

#### Tissue collection

All male mice, unless stated otherwise, were sacrificed by cervical dislocation at the age of 18- 20 weeks in the second day of darkness at the indicated circadian times (CT), which is considered as the same indicated time points used for normal LD. This was performed independent of external timing cues (Zeitgeber), such as the light-dark cycle to demonstrate rhythms generated by endogenous intestinal clocks ^78^. Eyes were removed prior to tissue dissection in dim red light. Tissues were harvested and directly transferred into RNA stabilization solution (NucleoProtect^®^ RNA, MACHEREY-NAGEL) overnight at 4 degrees and then stored in -80 degrees. For fixation, tissues were freshly harvested and then transferred into 4% Formaldehyde.

### Organoid

Freshly isolated tissue pieces from small intestine and colon were placed in phosphate-buffered saline (PBS) and 120µl 0.5M EDTA to detach the villi from the tissue. Tissues were changed to new tubes after 30 min incubation at 4 °C with gently shaking. Intestinal crypts were resuspended in Matrigel™ (BD Biosciences) and then seeded in 25 μl drops in 64-well plates after filtering and centrifuging. Plates were incubated at 37°C for 15min to allow suspension to polymerize before fresh medium was supplied. Organoid medium (IntesticultTM, Stemcell) was replaced every 3-4 days. Organoids for each time point were plated into a separate plate to limit manipulation or exposure to possible resetting cues, such as temperature. After synchronization with serum shock (50% Fetal Bovine Serum and 50% Intesticult) for 2 hours, medium was replaced with basic organoid medium (IntesticultTM, Stemcell). Every 6 hours over a 24-hour period (starting at 12h after serum shock) medium was removed and organoids were transferred to -80°C until further processing.

#### Gene expression analysis

RNA was extracted according to the manufacturer’s instructions (NucleoSpin^®^ RNA, MACHEREY-NAGEL) and measured by NanoPhotometer^®^N60 (IMPLEN). cDNA was synthesized from 1000ng RNA using cDNA synthesis kit Multiscribe RT (Thermofischer Scientific). qPCR was performed in a Light Cylcer 480 system (Roche Diagnostiscs, Mannheim, Germany) using Universal Probe Library system according to manufacturer’s instructions. Calculations (2^−ΔΔCt^ method) were normalized to elongation factor 1 alpha as housekeeper. For genes expression the following primers and probes were used: Brain and Muscle ARNT-Like 1 *(Bmal1)* F 5’-ATTCCAGGGGGAACCAGA-3’ R 5’-GGCGATGACCCTCTTATCC-3’ Probe 15, Period 2 *(Per2)* F 5’-TCCGAGTATATCGTGAAGAACG-3’ R 5’- CAGGATCTTCCCAGAAACCA-3’ probe 5, Nuclear receptor subfamily 1 group D member 1 *(Reverbα)* F 5’- AGGAGCTGGGCCTATTCAC-3’ R 5’-CGGTTCTTCAGCACCAGAG-3’ probe 1, D site-binding protein *(Dbp)* F 5’-ACAGCAAGCCCAAAGAACC-3’ R 5’- GAGGGCAGAGTTGCCTTG-3’ probe 94, Cryptochrome 1 (*Cry1*) F 5’- ATCGTGCGCATTTCACATAC-3’ R 5’- TCCGCCATTGAGTTCTATGAT-3’ probe 85, Tumor necrosis factor alpha *(Tnf)* F 5’- TGCCTATGTCTCAGCCTCTTC-3’ R 5’- GAGGCCATTTGGGAACTTCT-3’ probe 49, Interferon gamma (IFN-γ) F 5’- GGAGGAACTGGCAAAAGGAT-3’ R 5’- TTCAAGACTTCAAAGAGTCTGAGG-3’ probe 21, RNA abundance was normalized to the housekeeping gene Elongation factor 1-alpha *(Ef1a)* F 5’-GCCAAT TTCTGGTTGGAATG-3’ R 5’-GGTGACTTTCCATCCCTTGA-3’ probe 67.

### RNA sequencing

RNA quality was verified using an Agilent2100 Bioanalyzer with RNA 6000Nano Reagents. Library preparation and rRNA depletion was performed using the TruSeq Stranded mRNA Library Prep Kit. After the final QC, the libraries were sequenced in a paired-end mode (2x150 bases) in the Novaseq6000 sequencer (Illumina) with a depth of ≥ 12 Million paired reads per sample.

#### Pre-processing

The quality of Next Generation Sequencing data was assessed with FastQC v0.11.5 (RRID:SCR_014583, http://www.bioinformatics.babraham.ac.uk/projects/fastqc/). Adapter content and low quality reads were removed using Trimmomatic v0.39 (Anthony et al., 2014)

Trimmed FASTQ files were then mapped against the mouse mm10 genome with the STAR v2.7.5c aligner (Dobin et al., 2013). Format conversions were performed using samtools v1.3.1 (Li et al., 2009). The featureCounts program v1.4.6 (Liao et al., 2014) was used to count reads located within an exon, do not overlap multiple features, with a threshold of MAPQ > = 4 and are not chimeric.

#### Normalization and differentially expressed genes analysis

DESeq2 version 1.22.0 (RRID:SCR_015687 (Love et al., 2014) was used to normalize the read count matrix and perform differential expression analysis. Bioconductor package ‘‘biomaRt’’ version 2.38(RRID:SCR_002987 (Durinck et al., 2009)) was used to map MGI symbols to Ensembl gene IDs. To identify general DEGs between WT and KO mice, an initial model (Genotype + Time) was used with filtering FC>1.5, Benjamini-Hochberg-Adj.p<0.05 to identify differentially expressed genes (DEGs). Then a multi-model (Genotype + Period + Genotype::Period) was used to compare day vs night differences.

Gene Ontology biological process enrichment was performed using clusterProfiler (RRID: SCR_016884^79^, GO terms were significantly enriched with the q value<0.05. All genes expressed with minimum 1 count in any of the samples were used as the background universe. Redundant terms were removed manually.

#### Circadian analysis

The rhythmicity of oscillating transcripts was measured by JTK cycle (Hughes et al., 2010) through the MetaCycle R package^80^.With the setting: Period=24h and adj.p < 0.05, filtered genes were then defined as rhythmic. Package compareRhythms^81^ was modified and applied to compare rhythmicity differences in transcripts between control and *Bmal1*^IEC-/-^ mice as previously described^21^.

### High-Throughput 16S Ribosomal RNA (rRNA) Gene Amplicon Sequencing Analysis

Genomic DNA was isolated from snap-frozen fecal pellets and sequenced as previously described^21, 57^. Briefly, DNA NucleoSpin gDNA columns (Machery-Nagel, No. 740230.250) were used for DNA purification. After amplification of the V3-V4 region of the 16S rRNA gene, the multiplexed samples were sequenced on an Illumina HiSeq in paired-end mode (2x250 bp) using the Rapid v2 chemistry. Two negative controls, consisting of DNA stabilizer without stool, were used for every 45 samples to control for artifacts and insure reproducibility. High-Quality sequences of read counts > 5000 were used for 16s rRNA data analysis. Reads FASTQ files were consequently processed using an in-house developed NGSToolkit (Version Toolkit 3.5.2_64) based on USEARCH 11^82^. After trimming and FASTQ quality check, quality filtered reads were merged, deduplicated, clustered and a denoised clustering approach was applied to generate zOTUs^83^. Taxonomic assignment was performed with the EZBiocloud database^84^. Data was further analyzed with the R-based pipeline RHEA^85^. Phylogenetic trees are generated through the software MegaX^86^. Trees were visualized and annotated with the use of the online tool EvolView (http://www.evolgenius.info/evolview/)^87^.

### Targeted metabolite analyses

#### Sample preparation for targeted metabolite analyses

Approximately 20 mg of mouse cecal content was weighed extracted by bead beating (3 times of 20s 6 m/s with 30s breaks) with FastPrep-24 5G bead beating grinder (MP Biomedicals) supplied with a CoolPrep adapter. To measure BA and SCFAs, we used multiple reaction monitoring method and 3-NPH method respectively, as described previously^88^. Analyst 1.7 software (Sciex, Darmstadt, Germany) were used for data acquisition.

#### Targeted bile acid measurement

Targeted bile acid measurement was performed as previously descrived^21^. Briefly, measurement was performed using a QTRAP 5500 triple quadrupole mass spectrometer (Sciex, Darmstadt, Germany) coupled to an ExionLC AD (Sciex, Darmstadt, Germany) ultrahigh performance liquid chromatography system. The MS parameters and LC conditions were optimized using commercially available standards of endogenous bile acids and deuterated bile acids, for the simultaneous quantification of selected 44 analytes. Data acquisition and instrumental control were performed with Analyst 1.7 software (Sciex, Darmstadt, Germany) as previously described^88^.

#### Targeted short-chain fatty acid measurement

As previously described^21^, in brief, 40 µL of the cecal extract and 15 µL of isotopically labeled standards (ca 50 µM) were mixed with 20 µL 120 mM EDC HCl-6% pyridine-solution and 20 µL of 200 mM 3-NPH HCL solution. The measurement system was the same as described above. Data acquisition and instrumental control were performed with Analyst 1.7 software (Sciex, Darmstadt, Germany).

### Untargeted metabolite analyses

Fecal samples were collected from *IL-10*^-/-Sv129^ mice and their littermates wild types every 3 hours over the course of a 24h day. Samples were directly snap frozen and stored at -80 ◦C upon metabolite extraction. The untargeted analysis was performed using a Nexera UHPLC system (Shimadzu, Duisburg, Germany) coupled to a Q-TOF mass spectrometer (TripleTOF 6600, AB Sciex, Darmstadt, Germany). Separation of the fecal samples was performed either using a UPLC BEH Amide 2.1 × 100 mm, 1.7 µm analytic column (Waters, Eschborn, Germany) with a 400 µL/min flow rate or with a Kinetex XB18 2.1 x 100 mm, 1.7 µm (Phenomenex, Aschaffenburg, Germany) with a 300 µL/min flow rate. For the HILIC-separation the settings were as follows: The mobile phase was 5 mM ammonium acetate in water (eluent A) and 5 mM ammonium acetate in acetonitrile/water (95/5, v/v) (eluent B). The gradient profile was 100% B from 0 to 1.5 min, 60% B at 8 min and 20% B at 10 min to 11.5 min and 100% B at 12 to 15 min. For the reversed-phase separation eluent A was 0.1% formic acid and eluent B was 0.1% formic acid in acetonitrile. The gradient profile started with 0.2% B which was held for 0.5 min. Afterwards the concentration of eluent B was increased to 100% until 10 min which was held for 3.25 min. Afterward the column was equilibrated at starting conditions. A volume of 5 µL per sample was injected. The autosampler was cooled to 10 °C and the column oven heated to 40 °C. Every tenth run a quality control (QC) sample which was pooled from all samples was injected. The samples were measured in a randomized order and in the Information Dependent Acquisition (IDA) mode. MS settings in the positive mode were as follows: Gas 1 55, Gas 2 65, Curtain gas 35, Temperature 500 °C, Ion Spray Voltage 5500, declustering potential 80. The mass range of the TOF MS and MS/MS scans were 50–2000 m/z and the collision energy was ramped from 15–55 V. MS settings in the negative mode were as follows: Gas 1 55, Gas 2 65, Cur 35, Temperature 500 °C, Ion Spray Voltage –4500, declustering potential –80. The mass range of the TOF MS and MS/MS scans were 50–2000 m/z and the collision energy was ramped from –15–55 V.

The “msconvert” from ProteoWizard^89^ were used to convert raw files to mzXML (de-noised by centroid peaks). The bioconductor/R package xcms^90^ was used for data processing and feature identification. More specifically, the matched filter algorithm was used to identify peaks (full width at half maximum set to 7.5 s). Then the peaks were grouped into features using the “peak density” method^90^. The area under the peaks was integrated to represent the abundance of features. The retention time was adjusted based on the peak groups presented in most of the samples. To annotate possible metabolites to identified features, the exact mass and MS2 fragmentation pattern of the measured features were compared to the records in HMDB^91^ and the public MS/MS database in MSDIAL^92^, referred to as MS1 and MS2 annotation, respectively.

The QC samples were used to control and remove the potential batch effect, t-test was used to compare the features’ intensity between the groups.

The associated untargeted metabolomics data are available at https://massive.ucsd.edu/ProteoSAFe/dataset.jsp?task=ea2d927ea408493ea65d4fae9354ce20

### Dextran sulfate sodium(DSS) induced experimental colitis

14 weeks old male *Bmal1^IEC-/-^* mice and controls were released into constant darkness since experimental protocol started. Briefly, *Bmal1^IEC-/-^* and control mice were administered with 2% DSS dissolved in drinking water for 5 days, followed with normal water for 2 days. The onset of each mouse was calculated separately and sacrificed at circadian time (CT) 7 on day 7, when most of the immune cells are peaking in colon lamina propria .

### Transfer experiment

Cecal microbiota (collected at CT13) from either *Bmal1^IEC-/-^* and their controls, or severely inflamed *IL-10*^-/-BL^^6^ mice (n=4, mixture) were gavaged into germ-free *IL-10*^-/-BL^^6^ recipient mice, or germ-free *Bmal1^IEC-/-^* and their control recipient mice, respectively. 100µl of 7x10^6^ bacteria/µl were used for gavaging each mouse. Mice were weekly monitored for bodyweight changes and feces was sampled and stored in DNA stabilizer at week 5 after gavage. Mice were sacrificed on the second day of constant darkness at CT1 and CT13. All mice were kept in the gnotobiology facility in isolators equipped with HEPA-filters at 22 ± 1 °C with a 12-h light/dark cycle (lights on 5 am till 5 pm). Mice were single housed and had ad libitum access to autoclaved chow (V1124-300, Sniff Diets, Soest, Germany) and autoclaved water.

#### Immune cell isolation from the lamina propria

Freshly isolated intestinal tissues were flipped, rinsed and cut into 1 cm pieces. After 15 min incubation and shaking in DMEM with 20 µL of 1M DTT, tissue pieces were transferred into 37°C PBS containing 200 µL of 150mM EDTA for 10 min shaking. Jejunum tissue was then digested at 37°C for approximately 15 min in Thermoshake (Gerhardt) with 0.6 mg/ml type VIII collagenase (Sigma-Aldrich). Colonic tissue was digested at 37°C for approximately 25 min with 0.85 mg/ml type V collagenase (Sigma-Aldrich), 1.25 mg/ml collagenase D (Sigma-Aldrich), 10µl/ml Amphotericin (100x), 1 mg/mL Dispase II, and 10 U/µL DNase-V (Sigma). Following digestion, cells were passed through a 40 µm strainer and washed with PBS. Consequently, cells were fixed with 2% PFA for 20 min and washed, and stored in RPMI at 4 °C until further processing.

#### Flow cytometry measurement

For intracellular stainings, cells were permeabilized with 0.5% saponin and stained with conjugated antibodies at dilution 1/100-1/50 for 30 min. Namely, anti CD8-PE, anti CD3 - PerCP/Cy5.5, anti CD4-FITC, anti IL-17a -PE/cy7, anti INFy- APC.Surface stainings were performed using anti CD11c-PE, anti CD11b-APC/Cy7, anti F4/80-PE/cy7, anti Ly6G-APC conjugated antibodies at dilution 1/50 for 30 min. Cells were washed and resuspended and passed through Invitrogen™ Attune™ NxT Flow Cytometer. Analysis was performed using FlowJo v10.7.2.

#### Histology

Fixated and dehydrated tissue sections were cut into 5 μm thick slices and consequently stained according to the following steps: xylene/ 5 min, xylene/ 5 min, Ethanol 100%/ 5 min, Ethanol 100%/ 5 min, Ethanol 96%/ 2 min, Ethanol 96%/ 2 min, Ethanol 70%/ 2 min, Ethanol 70%/ 2 min, Water/ 30 s, hematoxylin/ 2 min, tap water/ 15 s, Scotts Tap Water/ 30 s, Water/ 30 s, Ethanol 96%/ 30 s, Eosin/ 30 s, Ethanol 96%/ 30s, Ethanol 96%/ 30 s, Ethanol 100%/ 30 s, Ethanol 100%/ 30 s, Xylene/ 90 s, Xylene/ 90s (Leica ST5020 multistainer). DPX new mounting media (Merck) was added to preserve the tissues. Histological scores were assessed blindly based on the degree of immune cell infiltration of all colonic wall layers (mucosa, submucosa and muscularis), crypt hyperplasia, goblet cell depletion and mucosal damage, resulting in a score from 0 (not inflamed) to 12 (severely inflamed) according to Katakura method (Katakura, Lee et al. 2005).

For AB/PAS staining, tissue slices were deparaffinized and rehydrated before being stained with Alcian blue solution for acidic mucins (1% volume/volume in 3% acetic acid, pH 2.5, 15 minutes), treated with periodic acid solution (0.5% volume/volume, 5 minutes) and co-stained with Schiff’s reagent for neutral mucins (Sigma-Aldrich, 10 minutes). Nuclei were then counterstained with hematoxylin. Consequently, tissue sections were differentiated (0.2% ammonia water), dehydrated, and mounted. The number of goblet cells was calculated as a total number per 100 μm2.

#### PICRUST 2.0

Sequence of the zOTUs which gained rhythmicity after restricted feeding in *IL-10*^-/-Sv129^ group were used to construct the metagenome using PICRUST2.0^33^ for prediction of functional of metagenomic functionality. Corrected zOTU 16 s rRNA gene copy number is multiplied by the predicted functionality to predicted the metagenome. Resulted enzymatic genes classified according to Enzyme Commission (EC) numbers were mapped to Metacyc pathways. Superclasses were removed and Metacyc pathways abundance was used for statistical analysis using STAMP (2.1.3). Statistical differences were calculated based on White’s non-parametric t-test and Benjamini Hochberg dales discovery rate to adjusted for multiple testing.

### Statistical Analyses

Statistical analyses were preformed using GraphPad Prism, version 9.0.0 (GraphPad Software), JTK_cycle v3.1.R^93^ or R. Between-sample microbiota diversity is calculated by generalized UniFrac using GUniFrac v1.1. distances within the Rhea^85^ pipeline. Microbiota composition comparison was calculated by PERMANOVA test on generalized Unifrac dissimilarity matrix and illustrated by MDS^85^. Circadian profile graphs, as well as phase calculation were analysed by fitting a cosine-wave equation: y=baseline + (amplitude·cos(2 · π · ((x-[phase shift)/24))). Non-parametric algorithm JTK_cycle was used for overall rhythmicity of all zOTUs and transcripts. Connected curves in circadian profile graphs within the figures indicate significant rhythmicity based on cosine analyses whereas connected straight lines indicate non-significant cosine fit. Comparison of rhythms between data sets was performed by adjusted version of compareRhythms as previously described^21^. Amplitude calculations depicted in the manhattan plots are based on the output of JTK_cycle and the phase was calculated by cosine-wave regression. Analysis between two groups was performed using the non-parametric Mann-Whitney test (two-sided). Two-way ANOVA followed with Benjamini-Hochberg correction was used to compare dataset with two categorical variables. Three-way ANOVA following Benjamini, Krieger and Yekutieli correction was used to compare dataset which contains three categorical factors. A p value ≤0.05 was assumed as statistically significant.

### Data availability

Data will be available from the Sequence Read Archive (SRA) upon request.

## SUPPLEMENTAL FIGURE LEGENDS

**Supplement Figure 1.**
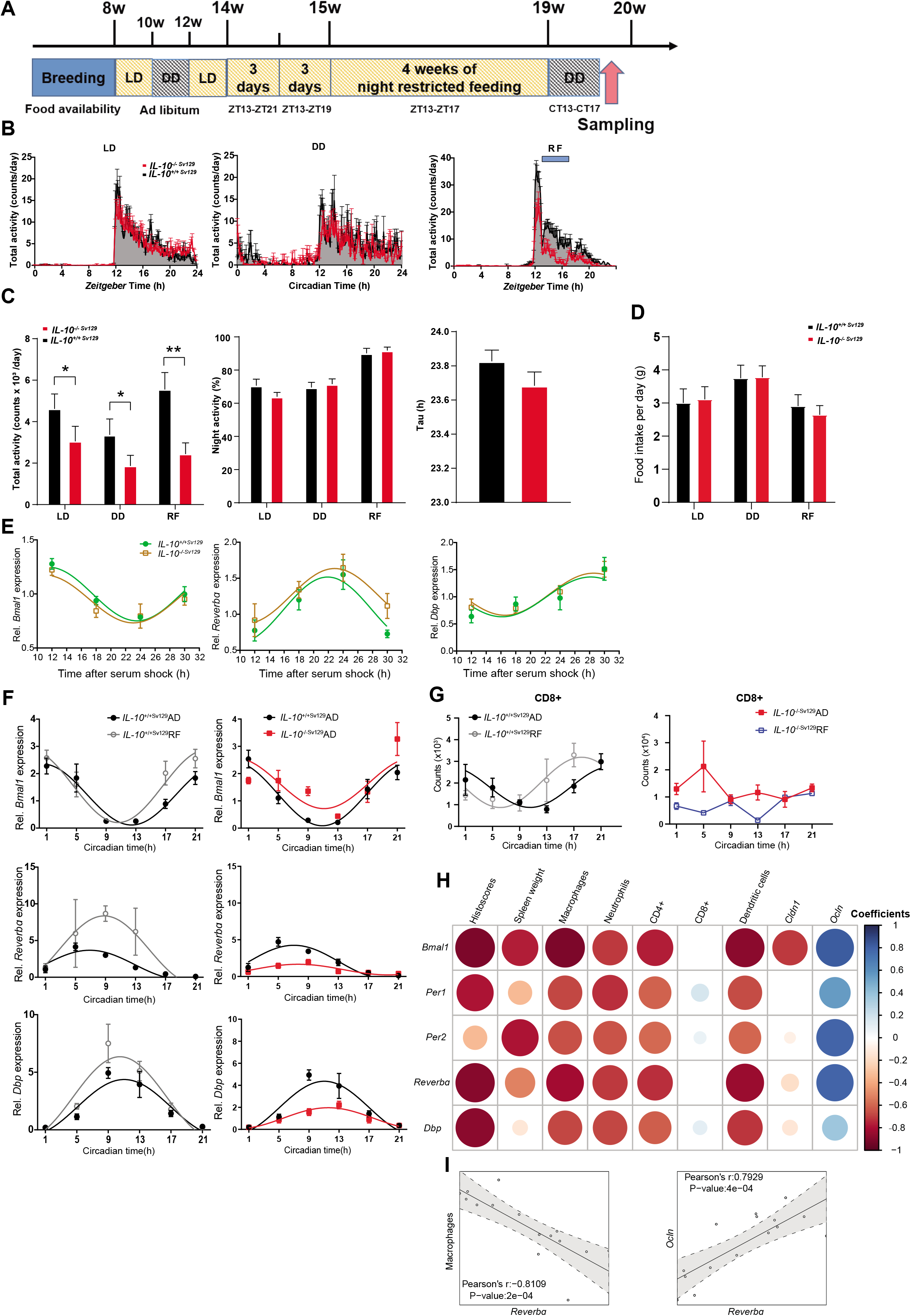
Circadian phenotype of IL-10^-/-Sv129^mice. (A) Schematic illustration of the experimental plan. (B) Representative graph of total activity of control and *IL-10^-/-^*^SV129^ mice for an individual day and (C) Quantification of total activity. (D) Food intake of control and *IL-10^-/-^*^SV129^ mice in LD, DD and RF. (E) Clock genes expression in jejunal organoids from control and *IL-10^-/-^*^SV129^ mice. (F) Colon clock genes expression in control (*IL-10*^+/+Sv129^) under AD and RF, and *IL-10^-/-^*^SV129^ mice under AD condition. (G) Circadian profile of the amount of CD8+ cells in colon lamina propria from control and *IL-10^-/-^*^SV129^ mice under AD and RF conditions. (H) Graphical display of all variables combinations in a matrix. Each correlation is depicted as a circle coloured according to the direction of correlation coefficients (negative, red; positive, blue). The size of the circles is dictated by the uncorrected p-value. (I) Representative correlation plot from (H). Significant rhythms are illustrated with fitted cosine-regression; data points connected by straight lines indicate no significant cosine fit curves (p > 0.05) and thus no rhythmicity. Significance were calculated by two-way ANOVA following Benjamini-Hochberg correction, * p ≤ 0.05, ** p ≤ 0.01, *** p ≤ 0.001, **** p ≤ 0.0001. Data are represented as mean ± SEM.

**Supplement Figure 2.**
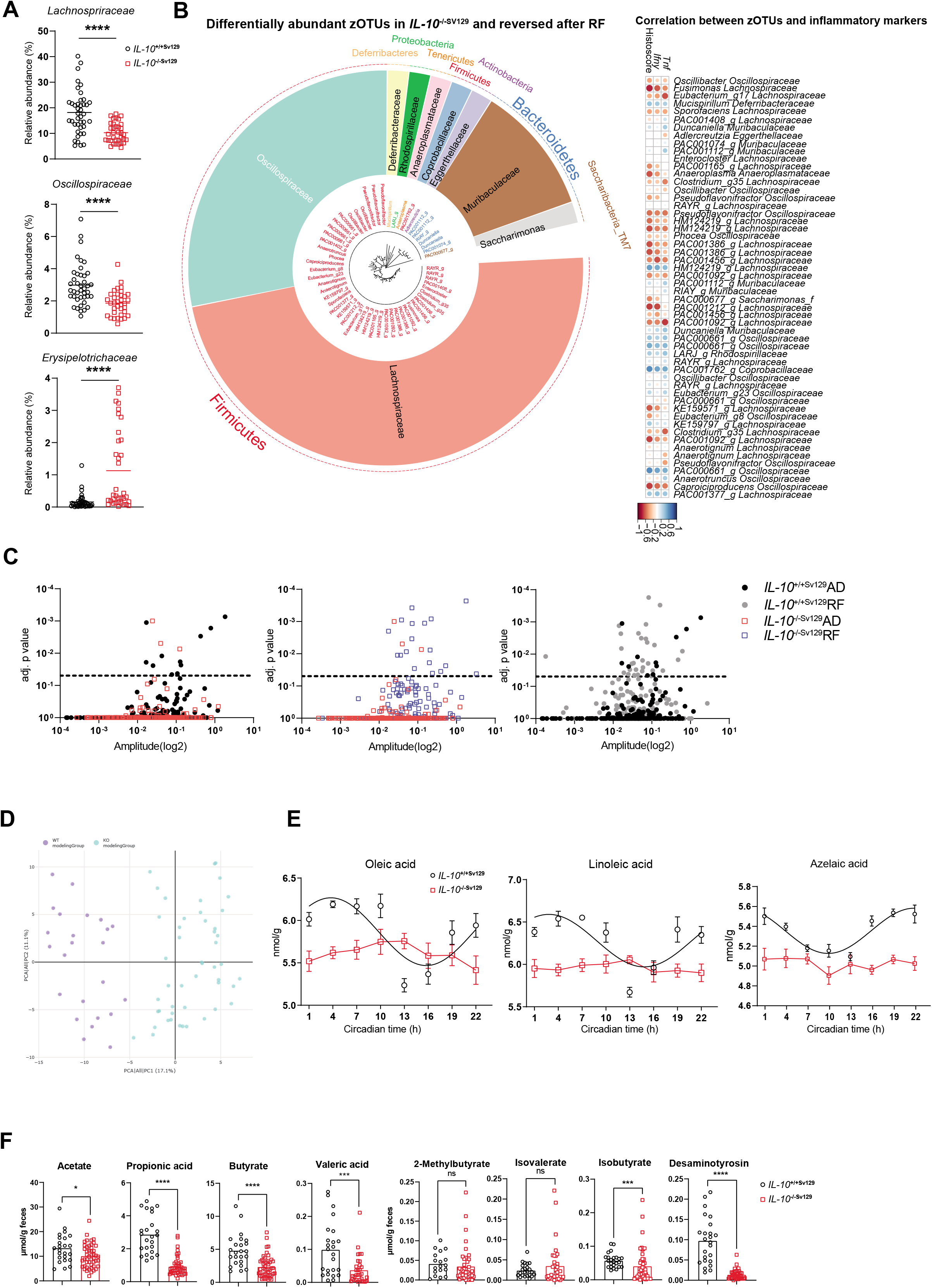
Alterations in the microbiome between IL-10^-/-Sv129^ mice and controls. (A) Representative taxa (*Lachnospriraceae, Oscillospiraceae and Erysipelotrichaceae*) which differentially abundant in fecal samples from *IL-10*^-/-Sv129^ mice. (B) Taxonomic tree of fecal microbiota which differentially abundant in *IL-10*^-/-SV129^ mice and reversed after RF. Taxonomic ranks are from phylum (outer dashed ring), family (inner ring highlighted) to genera (middle, color coded according to phylum) which are indicated by the individual branches (left) and correlations of zOTUs in (B) with histoscores, *Tnf* and *Ifn-y* gene expression (right). (C) Manhattan plot of the amplitude and adj.p value of zOTUs identified in control and *IL-10*^-/-Sv129^ mice under AD and RF conditions. (D) PCA-plot of fecal metabolites obtained from control and *IL-10*^-/-Sv129^ mice through untargeted metabolomics and (E) examples of metabolites which lost rhythmicity in *IL-10*^-/-Sv129^ mice. (F) Quantification of SCFA and desaminotyrosine measured by targeted metabolomics in fecal samples from control and *IL-10*^-/-Sv129^ mice. Significant rhythms are illustrated with fitted cosine-regression; data points connected by straight lines indicate no significant cosine fit curves (p > 0.05) and thus no rhythmicity. Data are represented as mean ± SEM. Statistics were performed by Mann–Whitney U test. Asterisks indicate significant differences *p<0.05, **p<0.01, ***p<0.001.

**Supplement Figure 3.**
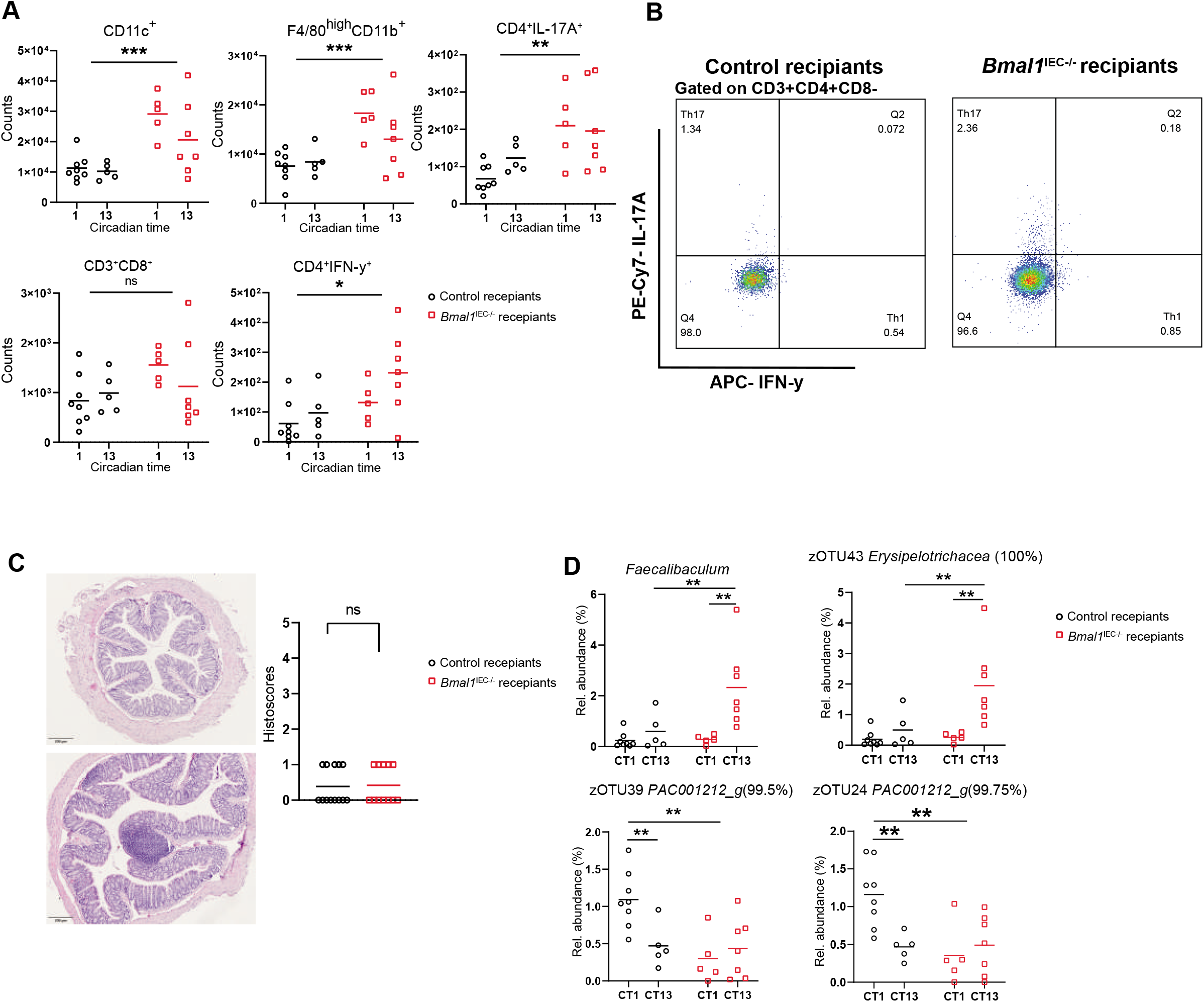
Disease-associated microbiota induce immune responses and microbial alterations in Germ-free Bmal1^IEC-/-^ mice. (A) Additional immune cells in the colon lamina propria from the recipients and (B) gating strategy for Th1 and Th17 cells. (C) Representative H&E staining scans of colon sections from *Bmal1^IEC-/-^* mice received control and disease-associated microbiota (left) and histopathological scores (right). (D) Representative microbial alteration in genus (*Faecalibaculum*) and specific zOTUs leves. Significance were calculated by two-way ANOVA following Benjamini-Hochberg correction, asterisks indicate significant differences *p<0.05, **p<0.01, ***p<0.001.

**Supplement Figure 4.**
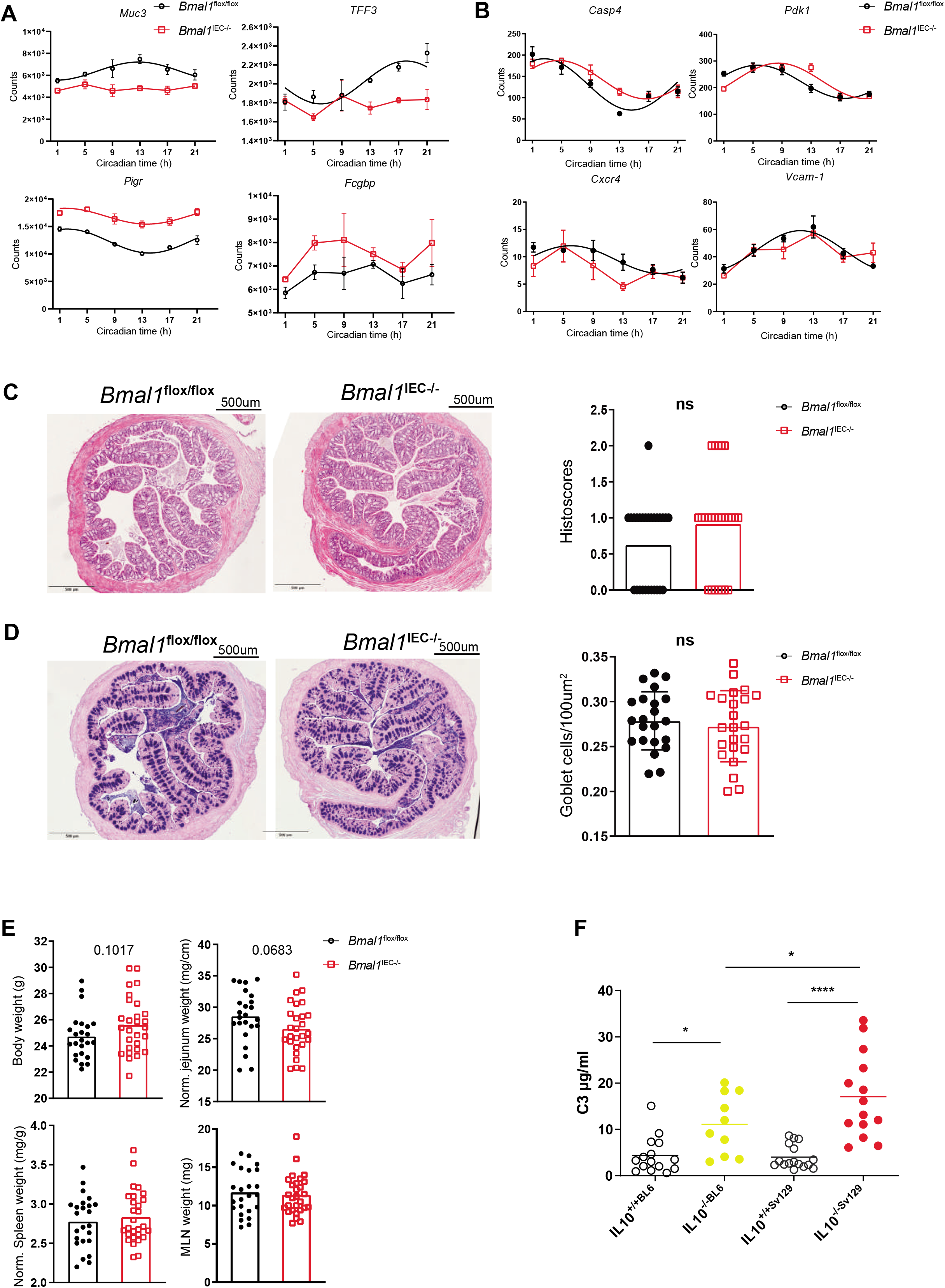
Inflammatory phenotype in the colon of Bmal1^IEC-/-^ mice. (A) Circadian profile of transcripts *Muc3, Tff3, Pig*r and *Fcgbp* and (B) *Casp4, Pdk1, Cxcr4* and *Vcam-1* in control and *Bmal1*^IEC-/-^ mice. (C) Representative scans of H&E staining of colon cross sections from control and *Bmal1*^IEC-/-^ mice (left) and histopathological scores (right). (D) Representative scans of PAS-AB staining of the same colon cross sections with (B) from control and *Bmal1*^IEC-/-^ mice (left) and quantification of goblet cells (right). (E) Body weight, organ weight of jejunum, spleen and MLN of control and *Bmal1*^IEC-/-^ mice. (E) Level of complement 3 in fecal samples from control and *IL-10*^-/-^ mice with BL6 and Sv129 background. Significant rhythms are illustrated with fitted cosine-regression; data points connected by straight lines indicate no significant cosine fit curves (p > 0.05) and thus no rhythmicity. Data are represented as mean ± SEM. Statistics were performed by Mann–Whitney U test and two-way ANOVA following Benjamini-Hochberg correction. Asterisks indicate significant differences *p<0.05, **p<0.01, ***p<0.001.

**Supplement Figure 5.**
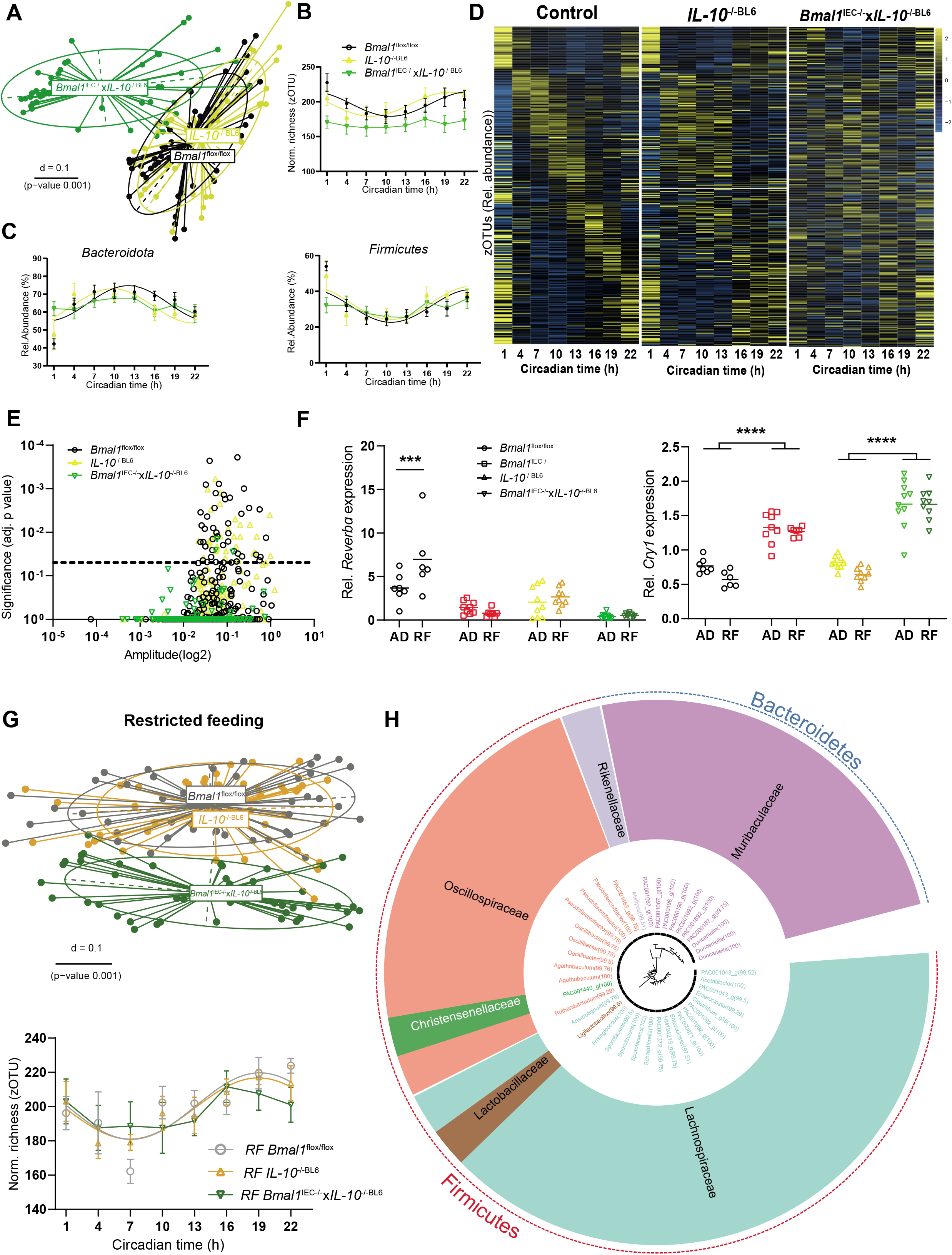
Disruption of microbial rhythms in Bmal1^IEC-/-^xIL-10^-/-^ mice. (A) Beta-diversity illustrated by MDS plots of fecal microbiota based on generalized UniFrac distances (GUniFrac) in control, *IL-10*^-/-Sv129^ and *Bmal1*^IEC-/-^x*IL-10*^-/-^ mice under AD condition. (B) Normalized richness and (C) abundance of major phyla (*Bacteroidota* and *Firmicutes*) in control, *IL-10*^-/-BL^^6^ and *Bmal1*^IEC-/-^x*IL-10*^-/-^ mice under AD condition. (D) Heatmap depicting the relative abundance of identified zOTUs (mean relative abundance>0.1%; prevalence > 10%). Data from control, *IL-10*^-/-BL^^6^ and *Bmal1*^IEC-/-^x*IL-10*^-/-^ mice are normalized to the peak of each zOTU and ordered by the peak phase in control group. Yellow means high abundance and blue low abundance. (E) Manhattan plot of the amplitude and adj.p value of zOTUs identified in control, *IL-10*^-/-BL^^6^ and *Bmal1*^IEC-/-^x*IL-10*^-/-^ mice. (F) Clock genes (*Reverba*, *Cry1)* expression measured at CT13 in colon of control, *IL-10*^-/-BL^^6^ and *Bmal1*^IEC-/-^x *IL-10*^-/-^ mice under AD and RF conditions. (G) Beta-diversity illustrated by MDS plots of fecal microbiota based on generalized UniFrac distances (GUniFrac) in control, *IL-10*^-/-Sv129^ and *Bmal1*^IEC-/-^x*IL- 10*^-/-^ mice (top) and normalized richness (bottom) under RF condition. (H) Taxonomic tree of fecal microbiota which have restored rhythms in *IL-10*^-/-^ mice after RF but not in *Bmal1*^IEC-/-^ x*IL-10*^-/-^ mice after RF. Taxonomic ranks are from phylum (outer dashed ring), family (inner ring highlighted) to genera (middle, color coded according to family) which are indicated by the individual branches. Significant rhythms are illustrated with fitted cosine-regression; data points connected by straight lines indicate no significant cosine fit curves (p > 0.05) and thus no rhythmicity. Data are represented as mean ± SEM. Statistics were performed by three-way ANOVA following Benjamini, Krieger and Yekutieli correction correction. Asterisks indicate significant differences *p<0.05, **p<0.01, ***p<0.001.

**Supplement table 1.**
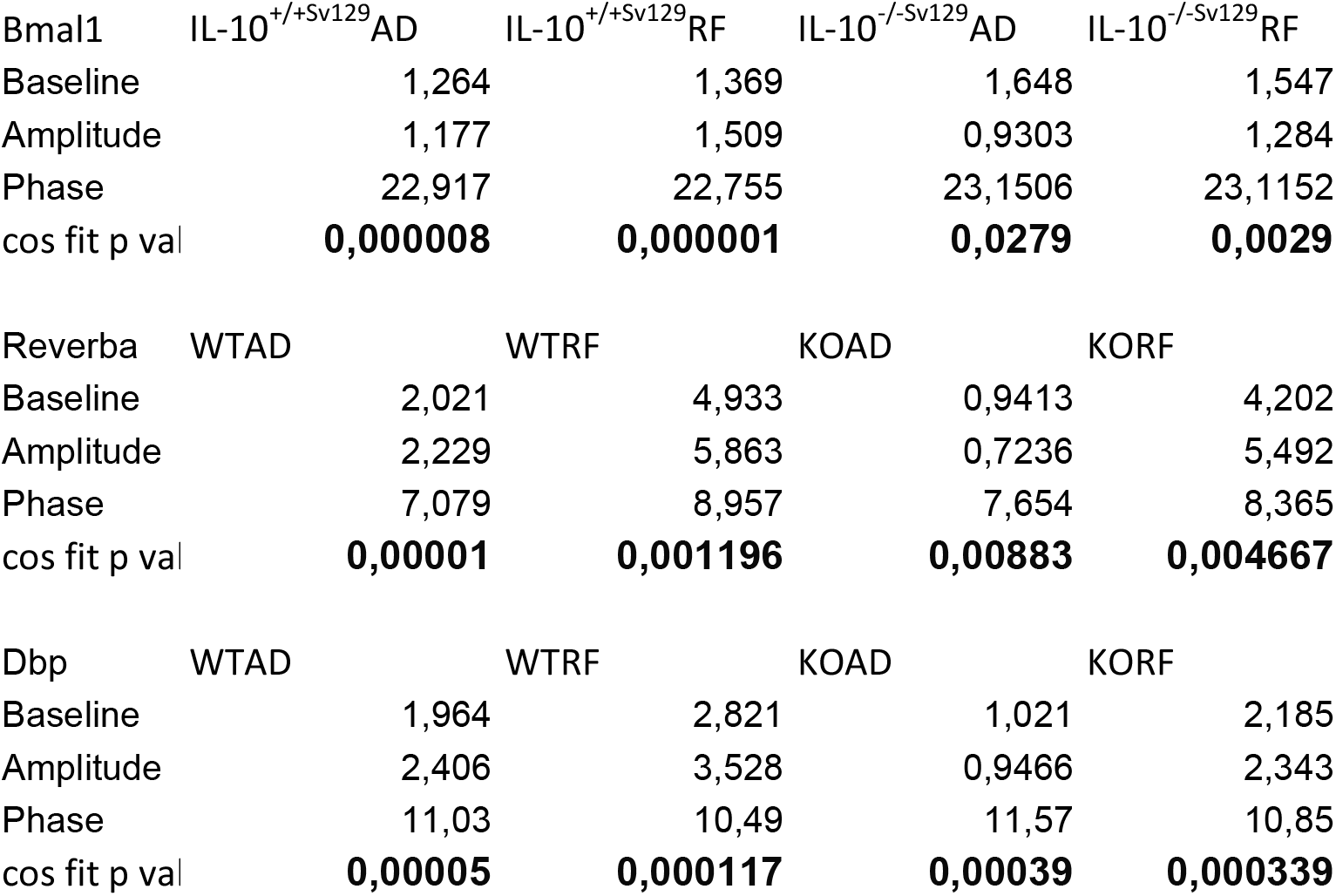
Circadian analysis of clock genes in IL-10^-/-Sv129^ mice under AD and RF.

